# Stratification of enterochromaffin cells by single-cell expression analysis

**DOI:** 10.1101/2023.08.24.554649

**Authors:** Yan Song, Linda J. Fothergill, Kari S. Lee, Brandon Y. Liu, Ada Koo, Mark Perelis, Shanti Diwakarla, Brid Callaghan, Jie Huang, Jill Wykosky, John B. Furness, Gene W. Yeo

**Affiliations:** Department of Cellular and Molecular Medicine, University of California San Diego, La Jolla, CA 92093, United States; Stem Cell Program, University of California San Diego, La Jolla, CA 92093, United States; Institute for Genomic Medicine, University of California San Diego, La Jolla, CA 92093, United States; Department of Anatomy & Physiology, University of Melbourne, Parkville, Victoria 3010, Australia; Florey Institute of Neuroscience and Mental Health, Parkville, Victoria 3010, Australia; Takeda Pharmaceuticals, San Diego, CA 92121, United States

**Keywords:** single cell transcriptome analysis, sensory, Piezo2, chemosensor, mechanosensor, enterochromaffin cells, enteroendocrine cells, 5-HT, serotonin, gut

## Abstract

Dynamic interactions between gut mucosal cells and the external environment are essential to maintain gut homeostasis. Enterochromaffin (EC) cells transduce both chemical and mechanical signals and produce 5-hydroxytryptamine (5-HT) to mediate disparate physiological responses. However, the molecular and cellular basis for functional diversity of ECs remains to be adequately defined. Here, we integrated single-cell transcriptomics with spatial image analysis to identify fourteen EC clusters that are topographically organized along the gut. Subtypes predicted to be sensitive to the chemical environment and mechanical forces were identified that express distinct transcription factors and hormones. A *Piezo2^+^* population in the distal colon was endowed with a distinctive neuronal signature. Using a combination of genetic, chemogenetic and pharmacological approaches, we demonstrated *Piezo2^+^* ECs are required for normal colon motility. Our study constructs a molecular map for ECs and offers a framework for deconvoluting EC cells with pleiotropic functions.

## INTRODUCTION

The capacity of the gut epithelium to sense and react to its surrounding environment is essential for proper homeostasis. Enteroendocrine (EEC) cells within the gut epithelium respond to a wide range of stimuli, such as dietary nutrients, irritants, microbiota products, and inflammatory agents by releasing a variety of hormones and neurotransmitters to relay sensory information to the nervous system, musculature, immune cells, and other tissues^1, 2^. In particular, enterochromaffin (EC) cells represent one of the major epithelial sensors. Historically, EC cells were histologically identified as the first type of gastrointestinal endocrine cells and have been thought of as a single cell type for about seven decades, until the emergence of recent studies that point to their heterogeneity^3–7^.

EC cells constitute less than 1% of the total intestinal epithelium cells, but they produce >90% of the body’s 5-hydroxytryptamine (5-HT, serotonin)^8^ to modulate a wide range of physiological functions^4–7^. Dysregulation of peripheral 5-HT levels is implicated in the pathogenesis of gastrointestinal (GI) diseases^9, 10^, cardiovascular disease^11^, osteoporosis^12^, and are associated with sudden infant death syndrome (SIDS)^13^ as well as psychiatric disorders, including autism spectrum disorders^14, 15^. The distinct and highly diverse functions of peripheral 5-HT suggest the possibility of specialization of EC sub-types that react to specific stimuli, such as chemicals in the lumen of the gut, mechanical forces, dietary toxins, microbiome metabolites, inflammatory mediators, and other GI hormones.

Since EC cells are infrequent and distributed throughout the gut wall, traditional approaches have utilized endocrine tumor cell lines, whole tissue preparations or genetic models (such as tryptophan hydroxylase 1 knockout) to investigate the functions of EC cells, which have generally assumed that EC cells are a single cell type and have not addressed their heterogeneity in sensory modalities. A recent study exploited intestinal organoids and described EC cells as polymodal chemical sensors, but lacked the resolution to disentangle the origin of the polymodality^16^. Some studies have compared small intestinal and colonic EC cells by RT-PCR^17, 18^, and one study used single-cell RNA sequencing (scRNA-seq) to compare a small number of human EC cells from the stomach and duodenum^19^. Single-cell transcriptomics have been utilized to profile intestinal epithelia^20, 21^ and enteroendocrine cells^22–24^, which include EC cells, however, questions regarding regional, molecular and functional heterogeneity of EC cells remain to be investigated in depth.

Here, we generated a genetic reporter of tryptophan hydroxylase 1 (Tph1), the rate-limiting enzyme of 5-HT synthesis in EC cells, and applied scRNA-seq to profile >6,000 EC cells. Together with spatial imaging analysis at single-cell resolution, we identified 14 clusters of EC cells distributed along the rostro-caudal and crypt-villus axes of the gut. We stratified EC subsets based on their repertoires of sensory molecules. In particular, we demonstrate an important role of the *Piezo2^+^/Ascl1^+^/Tph1^+^*subpopulation in normal gut motility, one of the proposed functions of EC-derived 5-HT^25–28^. Our comprehensive molecular resource and findings provide direct evidence of molecular and cellular heterogeneity of EC cells and is anticipated to be valuable for future studies.

## RESULTS

### Generation and characterization of a *Tph1-bacTRAP* reporter model

To systematically analyze EC cells we generated a *Tph1*-bacTRAP mouse strain by placing a *RPL10-GFP* fusion gene under the transcriptional control of the *Tph1* gene in a BAC construct (Supplementary Fig. 1a). In this *Tph1*-bacTRAP line, all GFP^+^ (representing Tph1^+^) cells in the duodenum and over 95% of the cells in the jejunum and large intestine were immunoreactive for 5-HT (Fig. 1a,b). Cells that stained positive for 5-HT but negative for GFP were also observed (Fig. 1b), which are likely to include tuft cells that store but do not synthesize 5-HT (Supplementary Fig. 1b)^29^. Epithelial cells from the duodenum, jejunum, ileum and colon were isolated from the *Tph1*-bacTRAP mice (Fig. 1c, and Supplementary Fig. 1c,d), sorted *via* fluorescence-activated cell sorting (FACS) and both GFP^+^ and GFP^-^ cells were subjected to scRNA-seq. Among a total of 4,729 cells, *Tph1* transcripts were measured in 88.9% of GFP^+^ cells, together with the chromogranin genes, *Chga* (in 97.3% of GFP^+^ cells) and *Chgb* (in 98.8% of GFP^+^ cells), established markers for EC cells (Supplementary Fig. 1e). Cluster analysis indicated that 23% of GFP^+^ cells were grouped with GFP^-^ cells (Supplementary Fig. 1d), although these cells had higher levels of the EC marker genes compared to the GFP^-^ cells they clustered with (Supplementary Fig. 1e,f). It is possible that some GFP is expressed in cells that have not yet fully committed to the EC lineage, or that there is some expression in cells outside this lineage, for example, in mast cells. Given the small sample size, we did not further investigate these cells in this dataset. In Supplementary Figures 1 d and f we refer to the GFP+ cells that clustered with the GFP-cells as “non-EC cells”.

**Figure 1.**
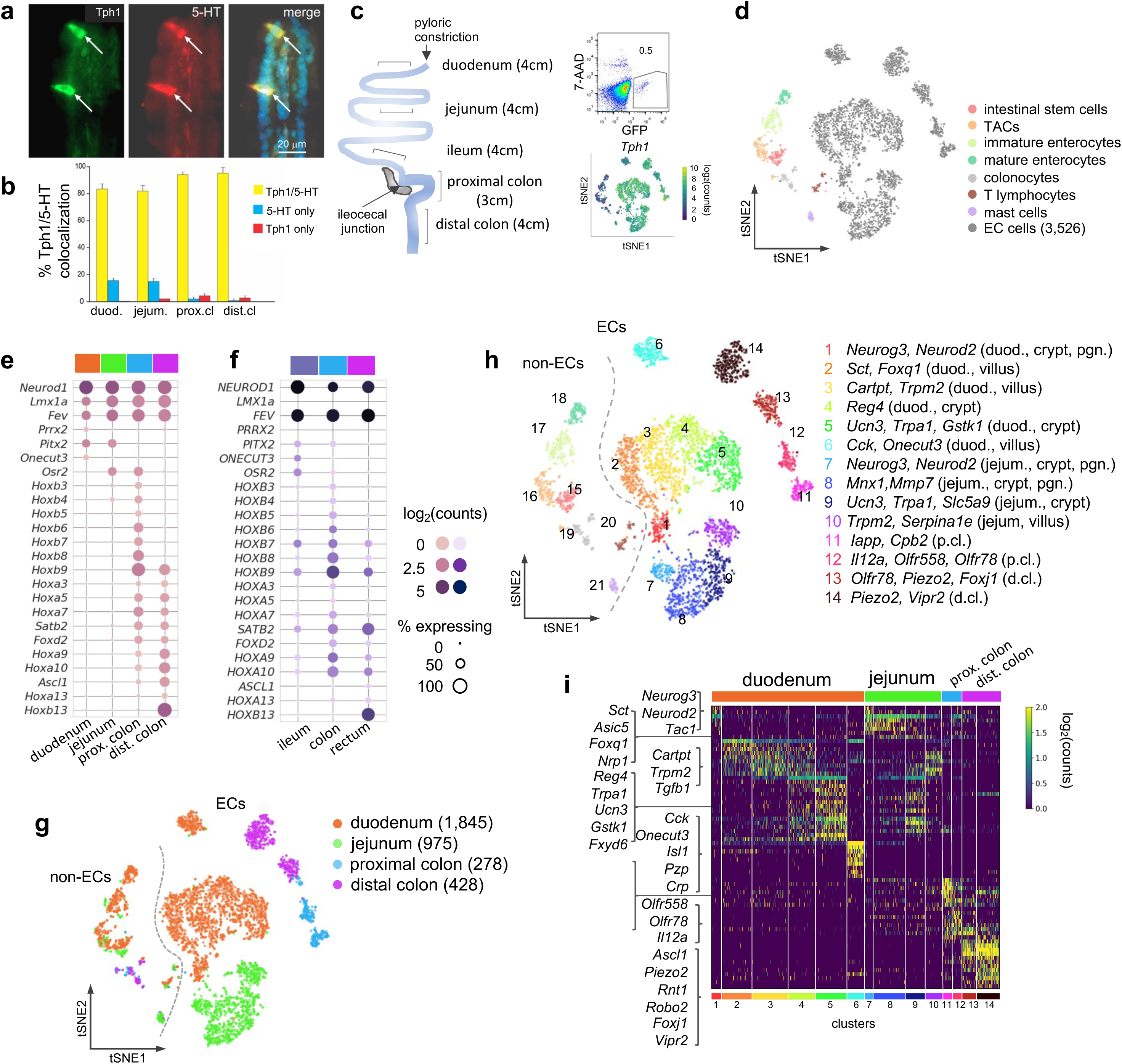
scRNA-seq identifies distinct intestinal enterochromaffin (EC) cell clusters. **(a,b)** Dual IF staining of 5-HT and GFP (representing Tph1) (a) and their quantification/colocalization in the indicated regions of *Tph1*-bacTRAP mice (b). **(c)** Schematic showing the isolation and enrichment of GFP^+^ cells from the indicated regions of the GI tract for scRNA-seq. *Upper right:* FACS plot of dissociated gut epithelial cells from *Tph1*-bacTRAP mice. Gate shows GFP^+^ cells, which account for ∼0.5% of total viable gut epithelial cells (7-AAD^-^ cells). *Lower right:* t-SNE projection of all GFP^+^ cells superimposed on an expression heatmap of *Tph1* in GFP+ cells. **(d)** t-SNE projection of all GFP^+^ cells isolated from *Tph1-bacTRAP* mice in the second cell profiling experiment, from which 3,526 enterochromaffin (EC) cells were identified and subjected to further analysis. TACs: transit amplifying cells; EC cells: enterochromaffin cells. **(e,f)** Transcription factors (TFs) that are differentially expressed in the mouse EC cells isolated from *Tph1*-bacTRAP mice along the rostro-caudal axis presented by regions as indicated in the color bar (e). Expression data of the human orthologues of the same TFs were extracted from human gut mucosa dataset (GSE125970). Enteroendocrine (EEC) cells were selected and presented by regions (f). Size of the circles represents percentage of expression and the intensity of the circles represents aggregated expression of indicated TFs in cells partitioned by regions. **(g)** t-SNE projection of all GFP^+^ cells color-coded by their regions in the GI tract. Numbers of cells retained from each region are indicated in parentheses. Dashed line demarcates the separation of non-EC cells (including stem cells, TACs, immature enterocytes, mature enterocytes, colonocytes, T lymphocytes and mast cells) versus EC cells. **(h)** tSNE projection of all GFP^+^ cells color-coded by clusters that were identified via the Louvain method. 14 clusters of EC cells (clusters 1-14) and 7 clusters of non-EC cells were identified (clusters 15-21, as described in (d). Key marker genes are listed for each cluster. duod.: duodenum, jejum.: jejunum, pgn: progenitor. **(i)** Heatmap of the top 5-10 signature genes for each cluster presented as normalized log_2_(counts) in all EC cells (in columns). Color-coded bar at the bottom represents the clusters identified in (h).

An independent single-cell profiling experiment was performed focusing on GFP^+^ cells (0.3-0.5% of total dissociated epithelial cells) from the duodenum, jejunum, proximal and distal colon of the *Tph1*-bacTRAP mice, where numbers of GFP^+^ cells were adequate (i.e. excluding the ileum) (Fig. 1d). 4,348 high-quality single cells were obtained, of which 19% comprised of non-EC cells and identified as stem cells, transit amplifying cells (TACs), immature enterocytes, mature enterocytes, colonocytes, T lymphocytes and mucosal mast cells based on their respective markers^20^ (Fig. 1d and Supplementary Fig. 1g). It is possible that the stem cell and transit amplifying cell clusters include cells that are in the process of differentiating into EC cells. However, given that they have not fully committed to the lineage, we do not consider it appropriate to classify them as “EC cells” for the purposes of analyzing EC cell types in this study. A total of 3,526 EC cells (at threshold >500 detected genes and <10% mitochondrial transcripts) were retained for analysis.

### Distinct EC subpopulations along the rostro-caudal axis

EC cells are one of the few EEC cell types distributed along the full length of the GI tract. The most significant transcriptomic distinction was observed between small intestinal and colonic EC cells as revealed by principal component analysis (PCA) (Supplementary Fig. 1h), even though all the EC cells expressed a core set of EC markers (Supplementary Fig. 1i). Unsupervised hierarchical clustering complemented with bootstrap resampling partitioned EC single cells by regions based on their overall transcriptomic similarity (Supplementary Fig. 1j). The regional distinction of EC cells is apparent from the examination of transcription factors (TFs) along the rostro-caudal axis. While *Pitx2* and *Osr2* demonstrated preferential enrichment in different segments of the gut, a suite of *Hox* genes were only observed in the colon, with *Hoxb13* specifically detected in the distal colon (Fig. 1e). This pattern was shared by all gut epithelial cells (Supplementary Fig. 1k) and was largely conserved in the human gut mucosa based on our comparative analysis of a scRNA-seq dataset of biopsy samples from human ileum, colon and rectum^21^. Notably, OSR2 and HOXB13 were preferentially enriched in the ileum and rectum respectively in these human samples (Fig. 1f). Consistent with a previous study, *Olfr78* and *Olfr558* were enriched in the colon but not the small intestine^18^.

To investigate the diversity of EC cells within each intestinal region, we clustered them using the Louvain method for unsupervised community detection and resolved 14 EC clusters that were mostly demarcated by regions: duodenum (clusters 1-6), jejunum (clusters 7-10), proximal colon (clusters 11-12), and distal colon (clusters 13-14, Fig. 1g,h). The EC cells from either duodenum (clusters 1-5) or jejunum (clusters 7-10) displayed a continuum in t-distributed stochastic neighbour embedding (tSNE) space, indicating a gradual transcriptomic change in the SI, whereas colonic EC cells formed clusters distinct from the SI clusters (Fig. 1g,h). Interestingly, cluster 6 (duodenum EC cells) was projected away from the rest of the SI EC cells in the tSNE space, suggesting a distinct molecular profile of the cells (see below).

### SI EC cells are predicted to switch sensors and hormone compositions along the crypt-villus axis

Next, we resolved the identities of the EC clusters using both known and newly identified marker genes (Fig. 1h,i). Since all intestinal EEC cells, including EC cells, derive from Neurog3^+^ cells, we annotated the *Neurog3*^+^ clusters as EC progenitors in the duodenum (cluster 1) and jejunum (cluster 7). Lineage tracing studies have established that crypt and villus EC cells preferentially express *Tac1* (encoding tachykinin precursor 1) and *Sct* (encoding secretin), respectively (independently validated in Supplementary Fig. 2b-e)^23, 30, 31^. We annotated clusters 4/5 (duodenum) and 8/9 (jejunum) as crypt clusters and clusters 2/3/6 (duodenum) and 10 (jejunum) as villus clusters (Fig. 2a) based on the relative expression levels of *Tac1* and *Sct*. We note that this division is not precise, but reflects a gradient between cells in the crypts and villi.

**Figure 2.**
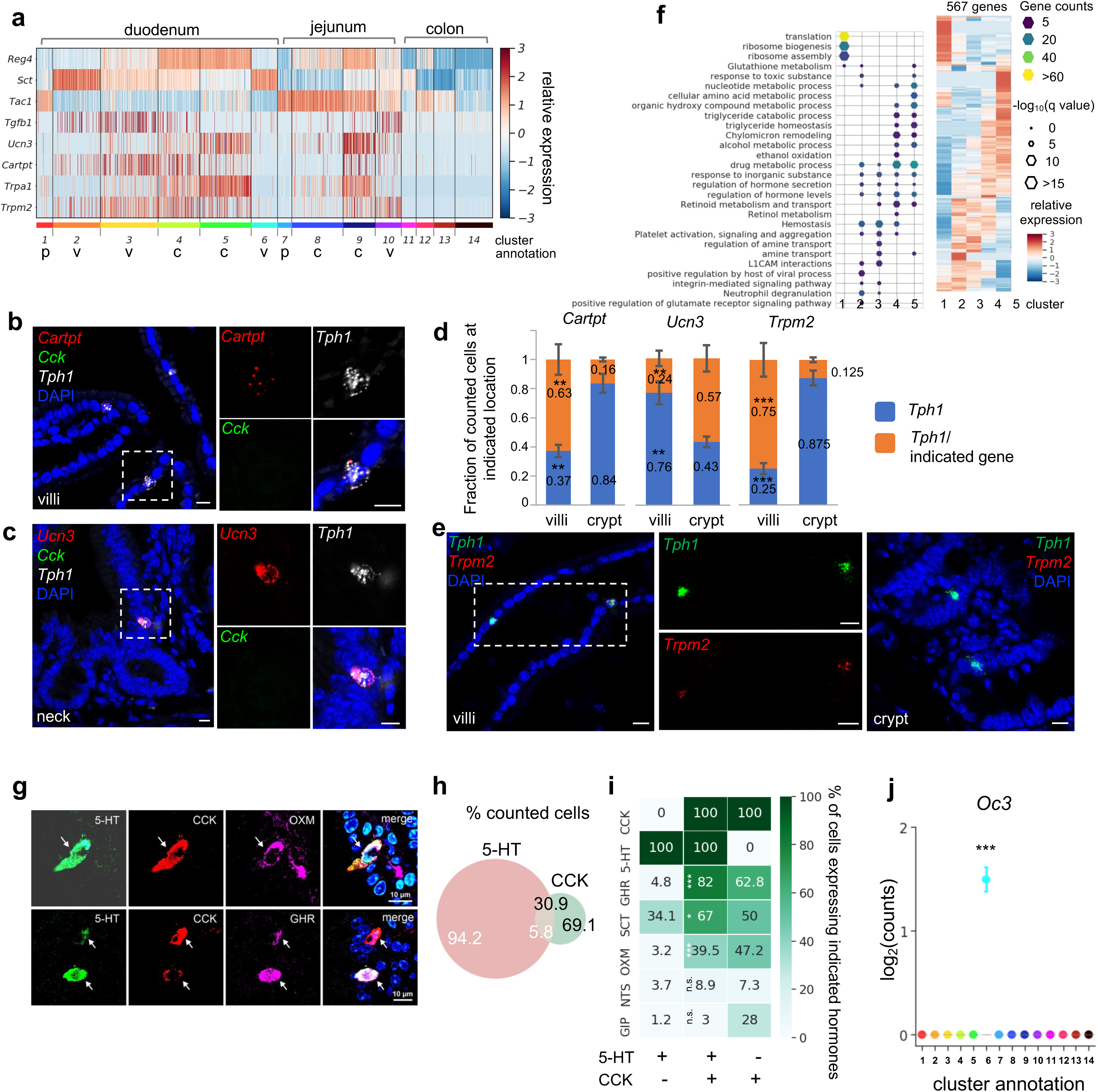
SI EC cells are predicted to switch sensors and hormone compositions along the crypt-villus axis. **(a)** Heatmap of representative genes with differential expression patterns between clusters annotated as crypt or villus. Relative gene expression (z-score) is shown across all single EC cells. Color-coded bar at the bottom represents the clusters. p: progenitor, v: villus, c: crypt. **(b,c,e)** smRNA-FISH of *Cartpt*, *Tph1*, and *Cck* (b); *Ucn3*, *Tph1*, and *Cck* (c); or *Trmp2* and *Tph1* (e). The boxed regions are enlarged on the right and split into individual channels. Data shown are representative examples from three independent mice. Scale bars: 10 µm. **(d)** Quantitation of *Cartpt*, *Ucn3* and *Trmp2* positivity in the *Tph1^+^* cells in the crypt versus villus. For each quantitation, 130-160 *Tph1*^+^ cells were counted from the crypt or villus of the duodenum in three independent mice. ** p<0.01, *** p<0.001; unpaired two-tailed Student’s *t*-test comparing the fractions in the villi against those in the crypt. **(f)** Gene ontology analysis of the 567DE genes identified in the duodenal clusters. DE genes were determined by FDR<10^−10^ against every other cluster (among clusters 1-5) based on Wilcoxon rank sum test and corrected by the Benjamini-Hochberg procedure. Relative expression (z-score) of DE genes is shown on the right, GO analysis of DE genes on the left. Size of the hexagons represents the q value of enrichment after -log_10_ transformation, and the density represents the number of genes per GO term. Accumulative hypergeometric testing was conducted for enrichment analysis and Sidak-Bonferroni correction was applied to correct for multiple testing. **(g)** Cells co-expressing 5-HT, CCK and a third hormone product shown by IF staining, such as a *Gcg* product (GHR), oxyntomodulin (OXM, upper panel) and the *Ghrl* product, ghrelin (GHR, lower panel). Arrows point to the triple-positive cells. Note that the relative levels of hormones vary considerably among cells, as shown in the 5-HT, CCK, ghrelin triple stain. **(h)** Venn diagram showing co-expression of 5-HT and CCK based on IF staining with indicated antibodies. Numbers represent percentages out of all 5-HT^+^ cells (white) or CCK^+^ cells (black). **(i)** Summary of 5-HT or CCK single-positive cells and 5-HT/CCK double-positive cells (by columns) producing a third hormone (by rows) as identified by IF staining. Heatmap and annotated numbers represent the percentage out of all cells in individual columns. ***p<10^−10^, *p<10^−2^ by hypergeometric tests against the 5-HT single-positive cells. (g-i) are based on IF staining for indicated peptides/hormones in three different animals. **(j)** Point plot depicting the median expression of *Oc3* in each cluster showing enrichment of *Oc3* expression in cluster 6 (*Cck^+^/Tph1^+^*cells) from the *Tph1-bacTRAP* dataset. Error bars represent upper and lower quantiles. *** p<10^−50^; two-tailed Kolmogorov-Smirnov statistic between the observed Oc3 distribution versus the bootstrap-facilitated randomization control distribution (median of n=500 randomizations shown in grey boxplots).

Since a spatial transcriptomic zonation along the crypt-villus axis had been demonstrated in intestinal enterocytes^32^ and EEC cells^23^, we further investigated whether EC cells diversify their transcriptome along the same axis to be poised for various functions. Using differential expression (DE) analysis, we identified a set of signature genes that were preferentially enriched in EC clusters annotated as being from the villus or crypt (Fig. 2a, Supplementary Fig. 2f). Their differential distribution was further tested to be statistically significant by comparing to bootstrap-facilitated randomization of gene subsets (Supplementary Fig. 2g,h).

To validate the spatial distribution of candidate genes, we utilized single-molecule RNA-FISH (smRNA-FISH) as an orthogonal method. We found that molecular sensors, such as *Trpa1* and *Trpm2*, together with additional hormone peptides, such as *Ucn3* and *Cartpt*, were differentially distributed in the EC cells along the crypt-villus axis. To illustrate, scRNA-seq suggested that *Ucn3* (encoding urocortin3) was frequently observed in the crypt clusters 4/5, whereas *Cartpt* (encoding cocaine and amphetamine regulated transcript prepropeptide) was enriched in the villus cluster 3 and to a lesser degree in cluster 4 (Fig. 2a). Consistently, smRNA-FISH demonstrated that *Ucn3* and *Cartpt* were selectively co-expressed with *Tph1*, but not with *Cck* (encoding cholecystokinin; expressed in cluster 6 as discussed below) (Fig. 2b,c). Specifically, *Ucn3* was found in 57.0% (± 3.3%) of *Tph1^+^* cells at the crypt or the neck of the villus, but observed in 24.0% (±2.1%) of *Tph1^+^* at the villus. In contrast, *Cartpt* was found in 63% (±3.5%) and 16% (± 0.5%) of *Tph1^+^* cells in the villus and crypt, respectively (Fig. 2b-d). Another pair of examples is the phytochemical sensor *Trpa1* (encoding transient receptor potential cation channel, subfamily A, member 1) and a novel sensor gene *Trpm2* (encoding transient receptor potential cation channel, subfamily M, member 2). *Trpa1* was frequently detected in the crypt cluster 5 in scRNA-seq, in agreement with its crypt location previously reported in rodent and human intestine^33, 34^, whereas *Trpm2* cells were mostly distributed in the villus clusters based on scRNA-seq analysis and further validated by smRNA-FISH to be co-expressed with 75% (±4.1%) and 12.5% (±0.4%) of *Tph1^+^* cells in the villi and crypt, respectively (Fig. 2d,e). Taken together, our findings are suggestive of a concomitant signature switch in the EC cells from *Tac1/Unc3/Trpa1* to *Sct/Cartpt/Trpm2* as the cells migrate from the crypt to the villus. In addition, scRNA-seq indicated that genes associated with oxidative detoxification including peroxidases and oxygenases (e.g.*Gstk1*, *Alb, Fmo1 and Fmo2)* were preferentially enriched in the crypt clusters 5/4, in contrast to the villus clusters 2/3 (Supplementary Fig. 2f). Furthermore, *Reg4, Ucn3, Trpa1, Gstk1,* and *Fmo1* expression levels are very low in progenitor clusters 1 and 7, which is consistent with reports from a time-resolved lineage tracing model that suggest that these genes are expressed late in the differentiation process of EEC cells^23^.

To assess the functional states of the duodenal and jejunal EC clusters, we performed gene ontology (GO) enrichment analysis based on cluster-enriched genes (identified as log_2_(fold-change)>2, FDR<10^−10^) (Fig. 2f, data not shown). Consistent with its annotation as a progenitor cluster, cluster 1 was enriched with terms ‘translation’ and ‘ribosome biogenesis’, suggestive of a high protein production state. Crypt clusters 4/5 were enriched with terms related to metabolism and hydroxy compound/alcohol metabolic process/detoxification, whereas villus clusters 2/3 were enriched with terms ‘hemostasis’, ‘viral process’ and ‘neutrophil degranulation’, suggesting the villus clusters may be involved in host defense (also see below for cluster 6).

### *Cck, Oc3, and Tph1* identify an EC subpopulation with a dual sensory signature

Emerging data suggest considerable co-expression of hormones within individual EEC cells, including EC cells^1, 6, 23, 24^. Among all the EC clusters, the largest number of hormone-coding genes was observed in cluster 6 expressing the highest levels of *Sct, Cck* and *Ghrl*, followed by *Gcg* and *Nts* (Supplementary Fig. 2i). Cluster 6 was almost exclusively constituted of duodenal EC cells (198/203, 97.5%) and comprised 10.9% of all retained duodenal EC cells (Supplementary Fig. 3a), thus representing <0.1% of total duodenal epithelial cells. To validate this small population, we investigated the co-expression of hormonal products by immunohistochemistry in the duodenum and found that 5.8 (±1.4%) of 5-HT expressing cells were positive for CCK, representing 30.9 (±6.6%) of CCK positive cells (Fig. 2g-i). Notably, of the 5-HT/CCK double positive cells, a large proportion demonstrated positivity for ghrelin (GHR, 82%), oxyntomodulin (OXM, 39.5%), one of the peptide products of the pre-proglucagon gene (*Gcg*), neurotensin (NTS, 8.9%), and a small percentage was positive for glucose-dependent insulinotropic polypeptide (GIP, 3.0%). These percentages were significantly reduced in 5-HT^+^/CCK^−^ cells, which presented 4.8%, 3.2%, 3.7%, and 1.2% positivity for GHR, OXM, NTS and GIP, respectively (Fig 2i). Thus, cluster 6 represents a subpopulation of EC cells with a broader spectrum of hormones (referred to as *Cck^+^/Tph1^+^* hereafter).

In a survey of TFs^1, 23^ that potentially specify the *Cck^+^/Tph1^+^* cells, we found *Onecut3* (*Oc3*) to be highly enriched in cluster 6 cells (Fig. 2j and Supplementary Fig. 2a), which we validated in an independent scRNA-seq dataset of *Neurod1+* enteroendocrine cells (Fig. 3i,j and Supplementary Fig. 3h, see below). Using smRNA-FISH, we found that 100% of the *Cck^+^/Tph1^+^* cells were positive for *Oc3*, whereas only 11.4% (±2.1%) of *Cck^−^/Tph1^+^* cells and 58% (±3.7%) of *Cck^+^/Tph1^−^* cells were positive for *Oc3* (Fig. 3a-c). *Oc3* single-positive cells were rarely observed (1.8% ±2.4% of 970 counted cells examined for *Cck*, *Oc3*, and *Tph1*; Fig. 3b,d), indicating that *Oc3* is largely restricted to *Cck^+^* and/or *Tph1^+^* populations and may be associated with the specification of *Cck^+^/Tph1^+^*cells. Notably, triple-positive cells (*Cck^+^/Oc3^+^/Tph1^+^*) were more frequently observed in the villus (25 out of every 100 counted single-, double-, or triple-positive cells) than in the crypt (6 out of every 100 single-, double-, or triple-positive cells; Fig. 3d). Focusing on *Oc3^+^*cells identified *via* smRNA-FISH, we found that the fraction of *Cck^+^/Oc3^+^* cells decreased from 69.3% (±0.98%) in the crypt to 22.8% (±5.3%) in the villus, while *Tph1^+^/Oc3^+^* cells increased from 17.3% (±2.4%) in the crypt to 38.0% (±3.2%) in the villus (Fig. 3d), suggesting a likelihood that *Cck^+^/Oc3^+^*double positive cells gradually acquire the ability to generate *Tph1* transcripts during their migration to the villus. Supporting this, triple-positive cells (*Cck^+^/Oc3^+^/Tph1^+^*) were found to express TFs shared with other EEC cells, including *Etv1*, *Isl1* and *Arx*, but displayed lower levels of *Lmx1α* enriched in the rest of *Tph1+* cells^35^ (Supplementary Fig. 3h).

**Figure 3.**
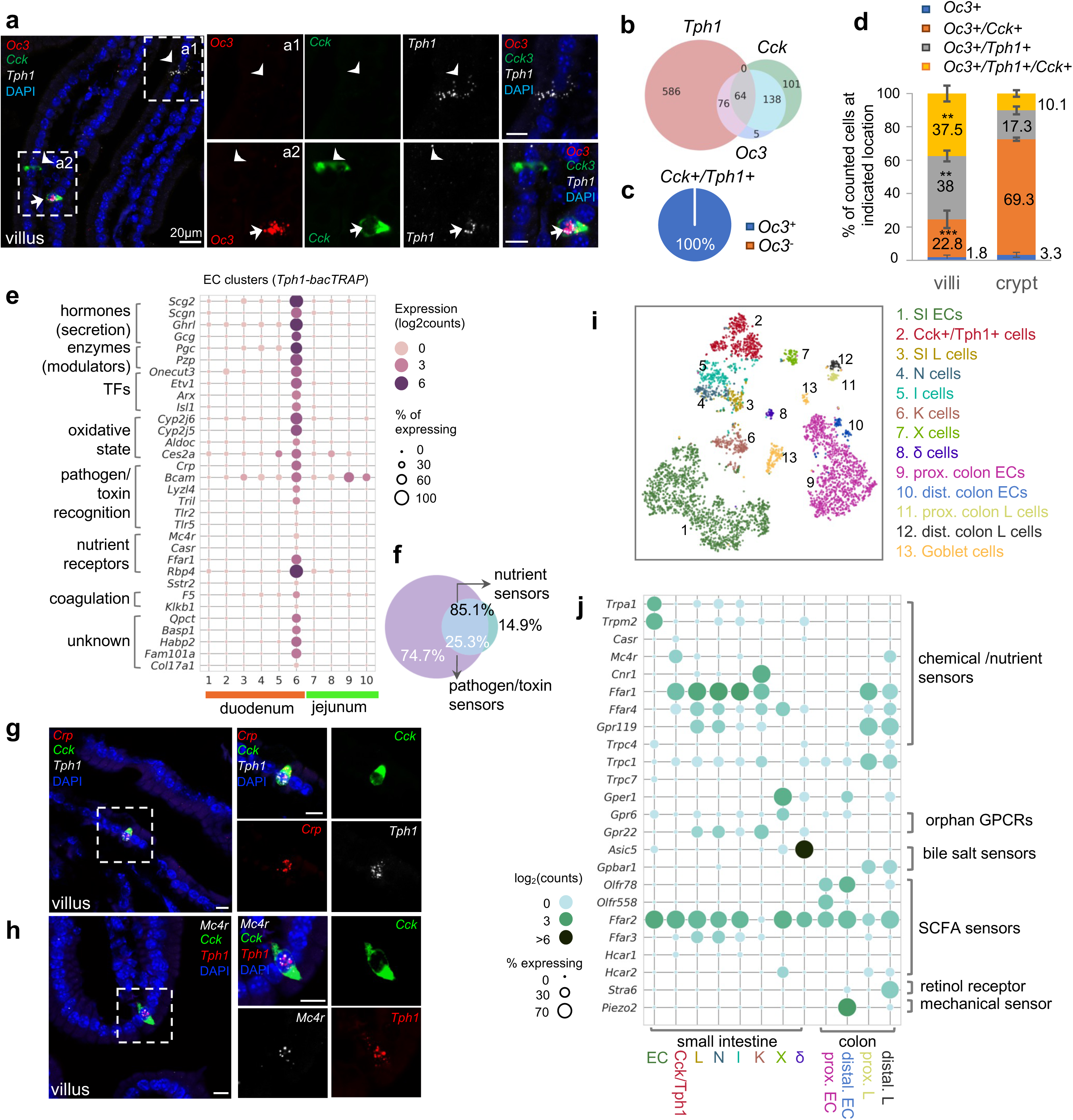
*Cck*, *Oc3* and *Tph1* specify an EC subpopulation with a dual sensory signature. **(a)** smRNA-FISH of *Cck/Oc3/Tph1.* Two boxed regions (a1 and a2) are enlarged on the right and split into individual channels. An arrow points to a *Cck^+^/Oc3^+^/Tph1^+^* cell in the villi and arrowheads point to *Tph1* or *Cck* single-positive cells with no *Oc3* expression in the villi. Images are representative from four different animals. Scale bars: 20 µm. **(b)** Venn diagram showing overlaps of *Tph1*, *Cck* and *Oc3*-expressing cells based on smRNA-FISH performed on duodenal sections. Numbers of counted cells are annotated based on four sections per mouse in four different animals. **(c)** Pie chart showing all counted *Cck^+^/Tph1^+^* cells partitioned by *Oc3* positivity based on smRNA-FISH. **(d)** Quantitation of *Oc3* expressing cells by smRNA-FISH in the villi versus in the crypts. ** p<0.01, *** p<0.001, unpaired two-tailed Student’s *t*-test comparing the percentages in the villi against those in the crypts. **(e)** Signature genes identified in *Cck^+^/Oc3^+^/Tph1^+^* cells by scRNA-seq in the *Tph1*-bacTRAP dataset. Size of the circles represents percentage of expression and intensity of the circles represents aggregated expression of indicated genes. **(f)** Venn diagram showing co-expression of two sets of genes summaried by the aggregated expression of genes associated with pathogen/toxin recognition (*Crp, Lyz4, Tril, Tlr2* and *Tlr5*) versus genes associated with nutrient sensing and homeostasis (*Mc4r* and *Casr*) in the *Tph1-bacTRAP* dataset. **(g,h)** smRNA-FISH of *Crp, Cck, and Tph1* (g) *and Mc4r, Cck, and Tph1*(h). The boxed regions are enlarged on the right and split into individual channels. Data are representative from three different animals. Scale bars: 10 µm. **(i)** tSNE representation of single *Neurod1-tdTomato^+^* cells isolated from the gut. Cells are color-coded by clusters that were identified via the Louvain method. **(j)** Dot plot of genes coding GPCRs, TRP channels, SLC transporters, purinergic receptors, and prostaglandin receptors, identified in clusters as shown in (i). Genes were selected if detected in >10% of cells in at least one of the indicated clusters and excluded if detected in >20% of enterocytes or colonocytes. Size of the circles represents percentage of expression and intensity of the circles represents aggregated expression of indicated genes partitioned by clusters.

We further determined the molecular signature unique to *Cck^+^/Oc3^+^/Tph1^+^* cells by contrasting them to the rest of the EC cells (Fig. 3e), or other EEC cells (Fig. 3j and Supplementary Fig. 3h,i). Specifically, discrete expression of *Crp* (encoding C-reactive protein) was identified in cluster 6 by scRNA-seq and mapped to *Cck+/Tph1+* cells by smRNA-FISH (Fig. 3g). Crp is a secreted bacterial pattern-recognition receptor involved in complement-mediated phagocytosis, known to be secreted by hepatocytes in response to inflammatory cytokines during infection or acute tissue injury^36^. Similarly, several genes encoding molecules recognizing pathogen-associated molecular patterns (PAMPs) were found in cluster 6 cells, including *Tril* (encoding TLR4 interactor with leucine rich repeats), *Tlr2* (encoding Toll-like receptor 2) and *Tlr5* (encoding Toll-like receptor 5). We validated the latter two by smRNA-FISH to be specific in *Cck^+^/Tph1^+^* cells (Supplementary Fig. 3b,c), supporting the notion that these cells play a role in pathogen or toxin recognition. Concordantly, *Bcam,* encoding a cell surface receptor that recognizes a major virulence factor (CNF1) of pathogenic *E. coli* ^37^, was enriched in cluster 6 (Fig. 3e). Along with cluster 6, the two villus clusters (cluster 2/3) were enriched with GO terms associated with host defense (Fig. 2f), suggesting a continuous evolution of EC cells along the crypt-villus axis to specify a complex subpopulation that mediates acute responses to tissue challenge. In addition to the unique signature of pathogen/toxin recognition, genes encoding G protein-coupled receptors (GPCRs) associated with nutrient sensing and energy homeostasis, *Mc4r* and *Casr*, were distinctly identified in cluster 6 cells (Fig. 3e). Mc4r (melanocortin receptor 4, encoded by *Mc4r*) plays a central role in energy homeostasis and satiety^38, 39^. CaR (calcium-sensing receptor, encoded by Casr) acts as a sensor for extracellular calcium and aromatic amino acids^40^. We further validated expression of *Mc4r* and *Casr* by smRNA-FISH to be specific to *Cck^+^/Tph1^+^* cells but not single-*Tph1^+^* or *Cck^+^* cells (Fig. 3h and Supplementary Fig. 3e,f). Although the nutrient sensing GPCRs were observed at relatively lower levels and frequencies in scRNA-seq data (Fig. 3e,j), among the *Mc4r^+^* or *Casr^+^* cells, >85% of them co-expressed at least one of the genes associated with pathogen/toxin recognition (Fig. 3f and Supplementary Fig. 3j). Together, these results suggest that a dual molecular signature associated with disparate functions can be resolved in the *Cck^+^/Tph1^+^/Oc3^+^*cells. Additionally, numerous genes encoding enzymes/enzyme modulators (*Pcg, Pzp, Habp2*) or proteins related with oxidation state (*Cyp2j5, Cyp2j6, Aldoc, etc*) were highly enriched in cluster 6 cells (Fig. 3e).

Finally, to provide a broader cellular context for the *Cck^+^/Oc3^+^/Tph1^+^*cells, we profiled *Neurod1^+^* EEC cells by scRNA-seq after crossing *Neurod1-Cre* mice with *Rosa26-LSL-tdTomato* mice. Among the 4,397 single EEC cells retained (at threshold >500 detected genes and <10% mitochondrial transcripts per cell) from duodenum, jejunum and colon, broad co-expression of hormone-coding transcripts was observed (Supplementary Fig. 3h) as in previous reports^20, 23^. To be compatible with conventional classification, clusters were annotated based on the most or second most abundant hormone-coding transcripts (Fig. 3i and Supplementary Fig. 3g,h). For simplicity, we assigned the diffuse clusters of *Tph1^+^* cells into either the SI EC cluster or proximal/distal colon EC clusters (based on *Hox* genes) and subdivided the *Cck^+^* cells into *Cck* dominating I cells, *Cck* and *Nts* co-expressing N cells, and *Cck* and *Gcg* co-expressing L cells, and the above described *Cck*^+^/*Tph1^+^*cells, reasoning that *Tph1* transcripts were otherwise restricted within the EC lineage (Supplementary Fig. 3g,h). The close relationship of I, N and L cells evident in our dataset is in agreement with a previous study revealing a temporal progression from L to I and N cells using a real-time EEC reporter mouse model^23^. In addition, *Zcchc12* and *Hhex* were found to be specifically expressed in X and δ clusters respectively, consistent with prior work^23^.

Among the *Neurod1+* EEC cells, a discrete cluster with high levels of *Cck*, *Sct*, *Tph1, and Ghrl*, together with *Gcg* and *Nts* transcripts, was identified and annotated as a *Cck^+^/Tph1^+^* cluster (Fig. 3i and Supplementary Fig. 3g,h). *Oc3* and key signature genes resolved in cluster 6 were also specifically identified in this *Cck^+^/Tph1^+^* population (Fig. 3j and Supplementary Fig. 3h,i). We thus conclude that the *Cck^+^/Tph1^+^* population in the *Neurod1-tdTomato* dataset is the equivalent of the cluster 6 in the *Tph1-GFP* dataset. A previous investigation with a *Neurog3* reporter identified *Oc3* in a subset of I and N cells, but did not resolve the molecular features of these cells^23^. Another example where these cells may have previously been identified is a single-cell analysis of proglucagon-expressing cells, which identified a cluster expressing *Tph1*, *Cck*, *Sct*, *Ghrl*, *Gcg*, and *Nts*, along with *Casr*, *Mc4r*, and *Pzp*^22^.

Taken together, our scRNA-seq profiling from two different genetic models, multiplex fluorescent smRNA-FISH and immunohistochemistry analysis of protein expression coordinately identified a discrete subpopulation of enteroendocrine cells preferentially located in the tip of villi in the duodenum and features a complex molecular signature, including a set of sensors associated with pathogen/toxin recognition and another set linked to nutrient sensing and homeostasis.

### Distinct molecular sensors are identified in EC cells versus other EEC cells

Having identified unique molecular sensors for various subpopulations of EC cells in the small intestine, we went on to evaluate whether these sensors are unique to EC cells by comparing them to other EECs. We focused on known and potential molecular sensors, including GPCRs, transient receptor potential channels (TRP channels), solute carrier transporters (SLCs), as well as purinergic receptors and prostaglandin receptors^41–43^ (Fig. 3j). In support of our previous findings, SI EC cells were preferentially enriched with *Trpa1* and *Trpm2* along with *Cartpt* and *Ucn3* transcripts. *Casr* was enriched in the *Cck^+^/Tph1^+^* cells, whereas *Mc4r* was primarily found in the *Cck^+^/Tph1^+^* cells from the SI and additionally detected in distal colonic L cells. Consistently with the findings from the *Tph1*-bacTRAP dataset, ∼99% of *Casr^+^* or *Mc4r^+^* cells in the *Cck^+^/Tph1^+^* cluster co-expressed at least one sensor gene associated with pathogen/toxin recognition (*Crp/Tlr2/Tlr5/Tril*) (Supplementary Fig. 3j), in support of our observation that *Cck^+^/Tph1^+^* cells are enriched with a dual sensory signature.

More broadly, in the EEC cells many sensors (*Cnr1, Asic5, Gper1* and *Stra6*) demonstrated a cluster-specific expression profile, while others (*Ffar1, Ffar4, Ffar2* and *Ffar3*) were widely distributed (Fig. 3j). In the SI, *Cnr1* (encoding CB1), *Asic5* (encoding a bile acid sensitive ion channel (Basic)^44, 45^, and *Gper1* (encoding G protein-coupled estrogen receptor 1, Gpr30) were found and validated by smRNA-FISH to be enriched in the *Gip*-dominant K cell^46^, delta cells^31^ and K cells, respectively (Supplementary Fig. 4a-d). In the colon, *Stra6,* (encoding a retinol transporter) was exclusively identified in distal colonic L cells and mapped to *Pyy^+^* cells by smRNA-FISH (Supplementary Fig. 4e,f). Transcripts encoding retinol binding proteins Rpb2 and Rpb4 have previously been identified in EECs, and have been implicated in cell differentiation processes^24, 47^. *Gpbar1* (encoding a bile acid sensor Tgr5) was found in the proximal/distal colonic L cells, rather than in the EC cells. We further validated this finding by smRNA-FISH (Supplementary Fig. 4g,h) and by a similar finding from human mucosa single cells (Supplementary Fig. 4i-k). However, our study may not have detected low levels of *Gpbar1* expression. A previous study suggested that *Gpbar1* is expressed in EC cells, but not enriched compared to other cell types^18^. In contrast to these cluster-specific sensors, the long chain fatty acid receptor *Ffar1* (*Gpr40*) was found in almost all *Cck^+^* cells, as well as in *Gcg^+^* L cells from both SI and colon. *Ffar2 (Gpr43)* and *Ffar3 (Gpr41/42)* encode GPCRs that recognize microbial metabolites, the former of which we widely observed in all EEC cells except K cells, whereas the latter we primarily detected in the closely related L,I and N cells (Fig. 3j). Lastly, almost all of these molecular sensors were confined to EEC cells, with little or no expression observed in other epithelial cell types^20^ (Supplementary Fig. 4i). Therefore, we have validated the specificity of EC sensors using an independent scRNA-seq dataset together with various public datasets^20–23^ and determined the enrichment of known and newly identified chemical/nutrient sensors in various types of EEC cells.

### Two distinct clusters are identified in proximal colonic EC cells

The EC cells in the colon present distinct transcriptomic profile from their counterparts in SI. In the proximal colon, EC cells encompassed two clusters: *Iapp^+^/Cpb2^+^/Serpine1^+^/Sct^+^*cluster 11 and *Il12a^+^/Olfr558^+^/Olfr78^+^/Tac1^+^* cluster 12 (Fig 4a). GO analysis indicated that proximal colonic EC cells were involved in coagulation regulation (Supplementary Fig 5a), which is supported by the selective expression of *Cpb2* and *Serpine1* (Fig 4a), encoding the two known plasmin inhibitors, thrombin-activatable fibrinolysis inhibitor (TAFI) and plasminogen activator inhibitor-1 (PAI-1), respectively. *Iapp* was also validated in a subset of proximal colonic EC cells (cluster 11) by smRNA-FISH, but not in distal colonic EC cells (Supplementary Fig 5b).

**Figure 4.**
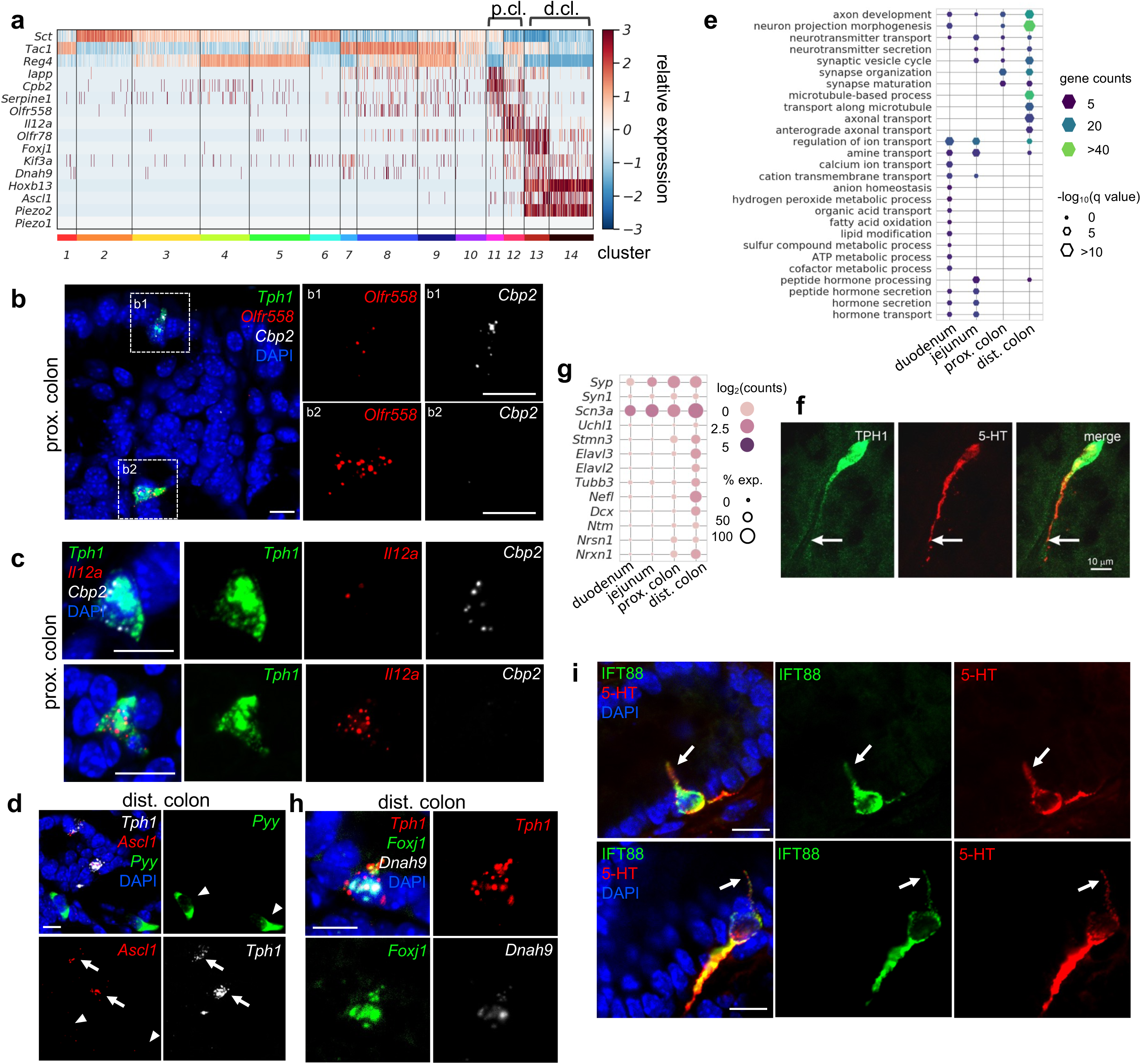
Distinct subpopulations of EC cells are resolved in the colon. **(a)** Heatmap showing the key signature genes identified in the colonic EC cells. Relative expression (z-score) of indicated genes are shown across all single EC cells based on the *Tph1-bacTRAP* dataset. Color bars at the bottom represent clusters and the ones on the top represent regions. p.cl.: proximal colon; d.cl.: distal colon. **(b)** smRNA-FISH of *Tph1, Olfr558*, and *Cpb2* in the proximal colon. Two dashed boxes (b1 and b2) are enlarged on the right to show two representative *Tph1^+^* cells with preferential *Cbp2* expression (b1) or *Olfr558* expression (b2). **(c)** smRNA-FISH of *Tph1*, *Il12a*, and *Cpb2* in the proximal colon. Two representative *Tph1^+^* cells with preferential *Cbp2* expression (upper) or *Il12a* expression (lower) are shown. **(d)** smRNA-FISH of *Ascl1*, *Tph1*, and *Pyy* in the distal colon. Arrows point to *Ascl1* staining in *Tph1^+^* cells. Arrowheads point to the absence of *Ascl1* staining in *Pyy^+^* cells. **(e)** GO analysis based on DE genes identified in regional EC cells. Hypothalamic neuron enriched genes were identified based on data from GEO dataset GSE74672, of which 60.7% genes were detected in gut EC cells. DE genes among the regional EC cells were determined by FDR<10^−10^ against every other region based on Wilcoxon rank sum test and corrected by Benjamini-Hochberg procedure. Size of the hexagons represents the q value of enrichment after -log_10_ transformation, and the density represents the number of genes per GO term. Accumulative hypergeometric testing was conducted for enrichment analysis and Sidak-Bonferroni correction was applied to correct for multiple testing. **(f)** IF staining against 5-HT and GFP (representing Tph1) in the distal colon. Note the prominent axon-like extension in the distal colon EC cell. **(g)** Dot plot showing representative neuron-related genes enriched in distal colon. **(h)** smRNA-FISH of *Tph1*, *Foxj1*, and *Dnah9* in the distal colon. **(i)** IF staining against IFT88 and 5-HT in the distal colon. Data shown are two representative examples from four mice. Scale bars in panels b-d,f,i: 10 µm. Images in panels b-d,f are representative from three mice.

Cluster 12 EC cells, on the other hand, are specialized in microorganism metabolite sensing. *Olfr558* (encoding olfactory receptor 558) and *Olfr78* (encoding olfactory receptor 78), two genes encoding GPCRs sensing short-chain fatty acids (SCFAs)^18, 48^, were enriched in cluster 12 (Fig. 4a,b). This is consistent with a previous report showing *Olfr78* and *Olfr558* in colonic EC cells with high expression levels of *Tac1*^24^. *Il12a* (encoding interleukin-12 subunit alpha) was concomitantly expressed in this cluster (Fig. 4a) and validated by smRNA-FISH (Fig 4c). *Il12a* is expressed by antigen-presenting cells, such as tissue-resident macrophages and dendritic cells, to promote helper T cells differentiation in responses to microbial infection. It is possible that cluster 12 EC cells may sense pathogens and transmit hormonal signals or (a) neurotransmitter(s) to evoke immune responses in the proximal colon. Using smRNA-FISH, we further mapped *Olfr558* and *Il12a* transcripts to a separate subset of EC cells expressing *Cpb2* (Fig. 4b,c), supporting the idea that there are subpopulations of EC cells in the proximal colon with gene transcripts associated with different physiological roles.

### Mechanosensor *Piezo2* is enriched in neuron-like distal colonic EC cells

Next, we focused on determining the unique properties of distal colonic EC cells. *Ascl1*, encoding Mash1 (mammalian achaete-scute homolog 1), is required for early neural tube specification^49–52^. Unexpectedly, *Ascl1* was found in the distal colonic EC cells (Fig. 1e, 4a; clusters 13 and 14) and was exclusively mapped to *Tph1^+^* cells but not to *Pyy^+^*cells in the distal colon, nor to any cell of the proximal colon or jejunum in smRNA-FISH analysis (Fig. 4d and Supplementary Fig 5c). Importantly, this feature was conserved in the human gut mucosa, where *ASCL1* was only detected in the *TPH1^+^*cells from the rectum (*HOXB13* high) (Supplementary Fig 5d).

Expression of *Ascl1* in distal colonic EC cells suggests a possibility these cells have acquired a neuronal-like profile. To test this, we compared EC cells with neuropeptidergic hypothalamic neurons^53^, which produce many hormone peptides similar to those found in the gut enteroendocrine cells. Prominently, EC cells from the distal colon, in contrast to their counterparts from other GI regions, were overwhelmingly associated with GO terms ‘neuron projection morphogenesis’, ‘axon development’ and ‘microtubule-based process’ (Fig. 4e and Supplementary Fig 5e). Numerous genes indicative of neuronal identity were mutually enriched in distal colonic ECs (Fig. 4g). Consistently, EC cells in the distal colon exhibited unique long basal processes, often extending for 50-100 µm, with 5-HT concentrated in the long processes, in contrast to the typical open-type, flask-shaped enteroendocrine cells observed in the proximal colon or SI (Fig. 4f and Supplementary Fig 5f)^54^. The function of these long basal processes is unknown but is unlikely to be related to synaptic transmission as there is no evidence of close apposition between the processes and neurons^54, 55^. Taken together, EC cells in the distal colon demonstrated molecular and cellular characteristics reminiscent of neurons, which is distinctive from all the other gut enteroendocrine cells.

Furthermore, we identified cilium-related features in a subset of the distal colonic EC cells. In cluster 13, *Foxj1* and *Dnah9*, encoding a TF (forkhead box protein J1) and an axonemal dynein (dynein heavy chain 9, axonemal) required for cilia formation^56, 57^, respectively, were selectively enriched (Fig. 4a) and validated by smRNA-FISH in the distal colonic EC cells but not in the proximal colonic EC cells or *Pyy^+^* L cells (Fig. 4h and Supplementary Fig. 5g). Genes associated with GO terms ‘cilium assembly/organization’ were also enriched in the distal colonic EC cells (Supplementary Fig. 5a,h). Primary cilium is a specialized cell surface projection that functions as a sensory organelle^58^, where GPCRs (e.g., rhodopsins, olfactory and taste receptors, Smo, *etc*) can be located to and sense the immediate surrounding environment to activate downstream signalling(s)^58, 59^. Consistent with the molecule features, we identified cilia by immunostaining against intraflagellar transport protein 88 (IFT88), an essential component for axonemal transportation, and found exclusive co-staining of IFT88 with 5-HT (Fig. 4i). Gene enrichment analysis of the *Foxj1^+^* EC cells also revealed concordant expression of *Olfr78* in cluster 13 (Fig. 4a), an observation validated by smRNA-FISH (Supplementary Fig. 5i) and suggesting that a subset of distal colonic EC cells (cluster 13) represent specialized sensory cells that detect microbial products.

Most prominently, in the distal colon, *Piezo2*, encoding a mechanosensitive ion channel, was identified in almost all EC cells (Fig. 4a). smRNA-FISH further revealed robust expression of *Piezo2* in the *Ascl1^+^/Tph1*^+^ cells residing in the epithelial layer of distal colon mucosa (Fig. 5a and Supplementary Fig. 6a). In addition, we noticed low levels (1 to 5 puncta per cell) of *Piezo2* signals in the lamina propria beneath the epithelium throughout the gut which contains mainly immune cells and connective tissue cells (Supplementary Fig. 6b). In contrast to *Piezo2*, *Piezo1* transcripts were either undetectable or sparsely observed (1 or 2 puncta per cell) in epithelium and lamina propria (Supplementary Fig. 6b). *Piezo2* was not detected in the *Tph1^+^* cells from the proximal colon, ileum, jejunum and duodenum *via* smRNA-FISH (Fig. 5b,c and Supplementary Fig. 6b). However, previous studies have demonstrated *Piezo2* expression in human and mouse small intestine by RT-PCR and confirmed localization in EC cells by immunohistochemistry, suggesting that *Piezo2* is expressed at levels below the detection threshold of the current study^60^. To further evaluate the variation of *Piezo2* expression, we sorted GFP^−^ and GFP^+^ cells from various segments of the gut isolated in the *Tph1-bacTRAP* animals and quantitated *Piezo2* and other newly identified signature genes by qPCR analysis (Fig. 5d and Supplementary Fig. 6c,d). A significant enrichment of *Piezo2*, up to 360-fold, was observed in the GFP^+^ cells sorted from the distal colon, when compared to the GFP^+^ cells from duodenum, jejunum, or proximal colon or to the GFP^−^ cells from the same regions. Based on multiple lines of evidence, we conclude that *Piezo2* is preferentially enriched in neuron-like distal colonic EC cells. Furthermore, a concomitant expression of *Piezo2* with *Foxj1* and *Olfr78* was revealed by scRNA-seq (Fig. 4a, cluster 13) and validated by smRNA-FISH (Supplementary Fig. 5i), which suggests that a subpopulation of *Piezo2^+^/Tph1^+^*cells are mechanosensory cells.

**Figure 5.**
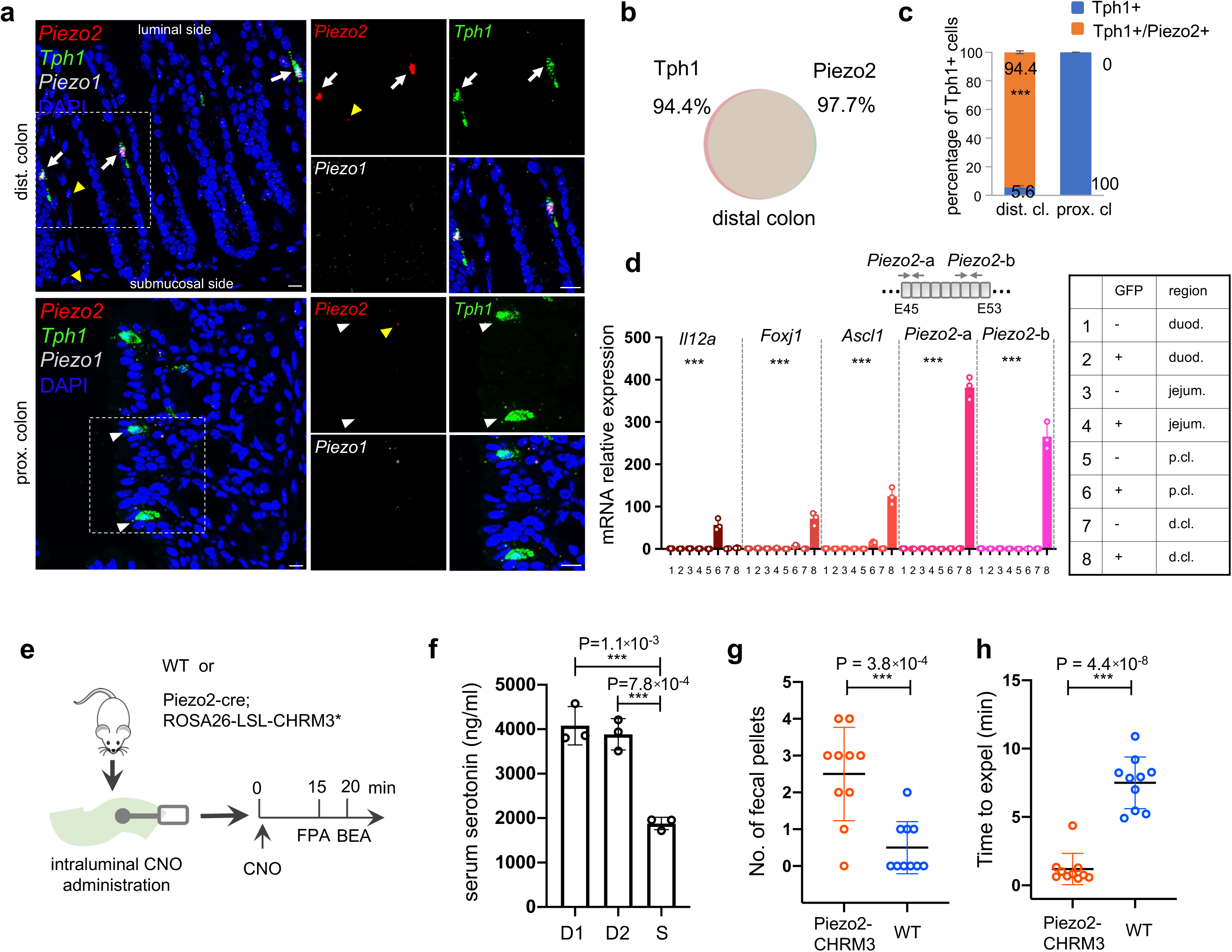
*Piezo2* is highly enriched in distal colonic EC cells that mediate colon motility. **(a)** smRNA-FISH of *Piezo2*, *Piezo1*, and *Tph1* in distal (*upper*) and proximal colon (*lower*). Arrows point to *Piezo2/Tph1* double-positive cells in the distal colon. White arrowheads point to absence of *Piezo2* in *Tph1^+^* cells in the proximal colon. Yellow arrowheads point to sparse staining of *Piezo2* within the lamina propria. Images are representative from four mice. Scale bars: 10 µm. **(b)** Venn diagram showing the co-expression of *Tph1* and *Piezo2* transcripts in the distal colon based on smRNA-FISH. Percentages are of double-positive cells out of respective single positive cells. A total of 274 cells were quantitated from four mice. **(c)** Quantitation of *Piezo2* and/or *Tph1* expressing cells in distal and proximal colon based on smRNA-FISH. A total of 386 cells were quantitated from four wild-type mice. *** p<0.001; unpaired two-tailed Student’*s t*-test of the double-positive fractions in distal colon versus proximal colon. **(d)** qPCR validation of regional enriched genes in EC cells. GFP^−^ and GFP^+^ cells were sorted from the duodenum, jejunum, proximal and distal colon in the *Tph1-bacTRAP* mice. Relative gene expression was computed relative to the values in the GFP^−^ cells isolated from the duodenum (indicated as 1), after normalization by the aggregates of three house-keeping genes (*B2m*, *Gapdh*, and *Rpl13a*). Two sets of primers were used to detect *Piezo2* expression. E45: exon 45 of the *Piezo2* gene (NCBI reference sequence NM_001039485.4). *** p<0.0001; two-way ANOVA test for both variables of GFP positivity and regions of gut, as well as the interaction of the two variables. Data are representative from three independent experiments. **(e)** Schematic of intraluminal administration of clozapine-N-oxide (CNO) followed by fecal pellet assay (FPA) and bead expulsion assay (BEA). FPA and BEA were conducted in the same cohorts at 15 and 20 minutes after CNO administration. **(f)** Serum serotonin level determined by ELISA from blood samples collected retro-orbitally 15 minutes after CNO administration (n = 3 mice per group). S: saline control, D1 (dose 1): 120 ng/kg, D2 (dose 2): 60 ng/kg. Representative data from two independent experiments are shown. Separate animal cohorts were used for serum serotonin assay versus FPA and BEA assays. **(g)** FPA from fecal pellets collected for 15 minutes after CNO administration. n = 10 in both WT and *Piezo2-CHRM3\** cohorts. Either WT or *Piezo2-cre;ROSA26-LSL-CHRM3\** (referred to as *Piezo2-CHRM3\**) mice were treated with CNO at 60ng/kg once intraluminally. Representative data from three independent experiments. **(h)** BEA performed 20 minutes after CNO administration. A glass bead was inserted 2 cm into the distal colon, and the time to expel it was monitored in each animal. n = 10 in both cohorts. Representative data from three independent experiments.

### *Piezo2^+^/Ascl1^+^/Tph1^+^* cells are required for normal colon motility

It is well documented that mechanical pressure or volume change within the gut lumen stimulates serotonin release from EC cells and initiates secretion and peristalsis^25, 26^. *Piezo2* is a mechanosensitive ion channel required in several pressure-sensing physiological systems^61–64^. A previous study demonstrated that the mouse jejunum and colon express functional mechanosensitive Piezo2 channels using *ex vivo* assays^65^. In addition, they found that mechanical stimulation evokes 5-HT release in primary colon cultures. A recent study has now shown that in an epithelial specific *Piezo2* knock-out model whole gut transit and colon transit times are slower compared to wild type animals and, in an *ex vivo* colonic motility assay, small shear forces increase frequency of colonic contractions in wild type but not knock out animals^66^. We have further validated the role of Piezo2 in colon motility, with a focus on the distal colon *Piezo2*^+^/*Ascl1^+^/Tph1^+^*cells.

First, we generated a *Piezo2-CHRM3\** model by crossing *Piezo2-IRES-cre* knock-in mice^63^ with *Cre*-dependent activating DREADD mice (*Rosa26-LSL-CHRM3*/mCitrine)*^67^, such that upon administration of clozapine-N-oxide (CNO), *Piezo2^+^* cells are chemically activated *in vivo* (Fig. 5e). 15 minutes after intracolonic administration of CNO, a robust elevation (2.1 ±0.2 fold) of serum serotonin was observed in *Piezo2-CHRM3\** mice, but not in WT, or *Rosa26-LSL-CHRM3\** mice following the same treatment, and no effect was observed with saline administration or in untreated animals (Fig. 5f and Supplementary Fig. 7a). To investigate serotonin release, we sorted mCitrine^+^/Epcam^+^ cells from various gut segments of the *Piezo2-CHRM3*/mCitrine* mice and evaluated serotonin release in response to CNO *in vitro* (Supplementary Fig. 7b-e). Despite the low levels of mCitrine signal, we identified and sorted ∼0.3% of mCitrine^+^/Epcam^+^ cells from the distal colon. In contrast, this population was largely absent from the proximal colon and duodenum, suggesting *Piezo2-cre* is primarily operational in the distal colon of the epithelial compartment. Additionally, total serotonin levels in the mCitrine^+^/Epcam^+^ cells from the distal colon were 4.7 (± 0.6) fold of those in the mCitrine^−^/Epcam^+^ cells from the distal colon, proximal colon and duodenum (Supplementary Fig. 7c). Moreover, in response to CNO stimulation, serotonin release from the mCitrine^+^/Epcam^+^ cells of distal colon was elevated by 1.9 (±0.19) fold, whereas mCitrine^−^/Epcam^+^ cells failed to respond (Supplementary Fig. 7d,e). Together, this data demonstrates that CNO-mediated activation of *Piezo2+* cells leads to robust serotonin release from epithelial EC cells.

We observed increased fecal pellet output from the *Piezo2-CHRM3\** mice within 15 minutes after CNO administration, in contrast to the WT controls (Fig. 5g). In a bead expulsion assay (BEA), colon motility was found to be significantly accelerated, such that the time to expel an inserted bead was shortened from 7.24 (±2.0) min in wild type controls to 1.16 (±1.17) min in the *Piezo2-CHRM3*/mCitrine* animals after CNO treatment, while no difference was observed between wild type controls and *Piezo2-CHRM3*/mCitrine* animals in untreated or saline treated cohorts (Fig. 5h and Supplementary Fig. 7f).

We noticed that *Htr4*, encoding the 5-HT4 receptor, a prokinetic 5-HT receptor when activated, was selectively expressed by epithelium in the deep crypts of distal colon (Supplementary Fig. 6e). Meanwhile, the long basal processes of EC cells filled with 5-HT always extend towards the base of the crypts, where the *Htr4* is preferentially expressed (Supplementary Fig. 6f). This finding suggests close proximity of 5-HT release to its receptor.

Next, to investigate whether *Piezo2^+^/Ascl1^+^/Tph1^+^* ECs are required for normal colon motility, we crossed *Piezo2-IRES-cre* with *Rosa26-LSL-DTR* (diphtheria toxin receptor)^68^ to generate *Piezo2-DTR* mice, such that upon diphtheria toxin (DT) administration, *Piezo2*^+^ cells would be depleted. Systemic administration of DT led to lethality in the *Piezo2-DTR* mice within 12 hours, but not in the *Rosa26-LSL-DTR* or *Piezo2-cre* mice (data not shown), likely due to the essential function of Piezo2 in respiration^62^. To avoid lethality, we administrated DT intraluminally into the distal colon for 5 consecutive days and assessed distal colon motility by a 2-hour fecal pellet assay (FPA) and the bead expulsion assay (BEA) (Fig. 6a). Profound co-depletion of *Piezo2* and *Tph1* transcripts was demonstrated by smRNA-FISH in the distal colon of the *Piezo2-DTR* mice, but not in the proximal colon or in the WT mice receiving the same DT treatment (Fig. 6b,c and Supplementary Fig. 8a). Substantial loss of 5-HT^+^ cells was further validated by immunofluorescence staining in the distal colon, but not in the proximal colon, while the general epithelial architecture was well-maintained (Fig. 6b,d and Supplementary Fig. 8a). Importantly, despite the extensive reduction of epithelial *Piezo2,* both the number and the intensity of the *Piezo2* puncta in the lamina propria of the *Piezo2-DTR* mice remained comparable to those of WT controls receiving the same DT treatment (Supplementary Fig. 8c-e), suggesting intraluminal administration of DT is unlikely to extensively perturb *Piezo2* expression in the lamina propria.

**Figure 6.**
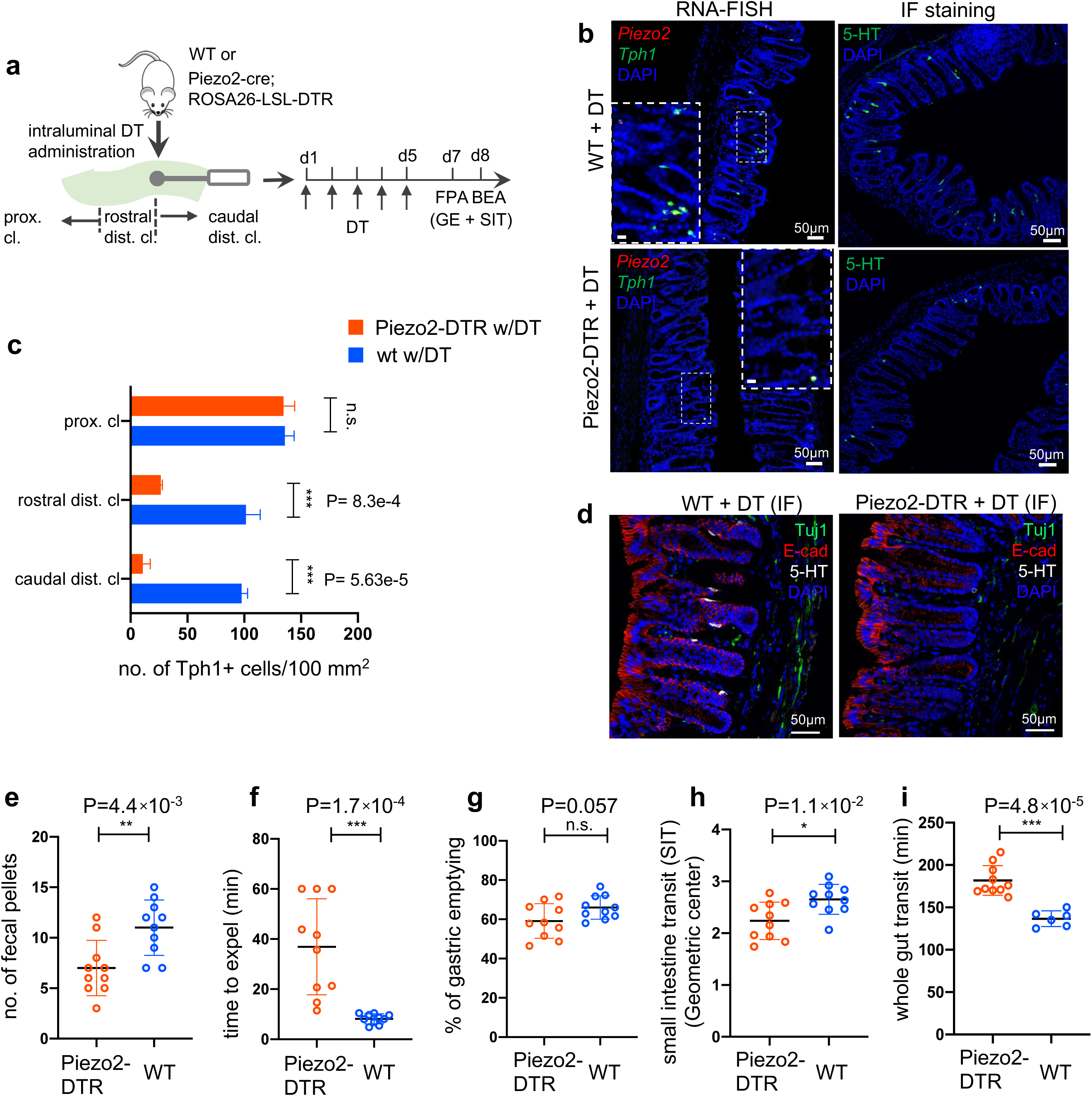
*Piezo2^+^/Ascl1^+^/Tph1^+^* cells are required for normal colon motility. **(a)** Schematic of intraluminal administration of diphtheria toxin (DT) followed by fecal pellet assay (FPA) and bead expulsion assay (BEA) in the same cohort. In a separate cohort, gastric emptying (GE) and small intestine transit (SIT) time assays were conducted following DT administration. Either wild type or *Piezo2-cre;ROSA26-LSL-DTR* (referred to as *Piezo2-DTR*) mice were treated with DT at 50 µg/kg twice a day for five consecutive days. **(b)** smRNA-FISH assay of *Piezo2* /*Tph1* (*left*) and IF staining for 5-HT (*right*) from the distal colon in wild type (*upper*) and *Piezo2-DTR* (*lower*) mice after DT administration. Dashed boxes are enlarged in the insets. Images are representative from five animals per group. Scale bars: 50 µm **(c)** *Tph1*^+^ cell counts based on smRNA-FISH experiments from the caudal distal colon (caudal dist. cl.), rostral distal colon (rostral dist. cl.), and the proximal colon (proximal cl.) for animals shown in (a) and (b). Five sections per animal were examined in five animals per group. **(d)** IF staining of 5-HT, E-cadherin and Tuj1 in the distal colon of WT (left) and *Piezo2-DTR* (right) mice after DT administration. Images are representative of five different animals per group. Scale bars: 50 µm **(e-f)** FPA showing the number of fecal pellets collected in 2 hours (e), and BEA measuring the time to expel a glass bead inserted 2 cm into the distal colon (f). n=10 per group, representative data from four independent experiments. **(g,h)** Gastric emptying time (g) and small intestine transit (SIT) time (h) examined after five days of consecutive treatment of DT. Animals were orally gavaged with methylcellulose supplemented with rhodamine B dextran (10 mg/ml). 15 minutes after gavage, the remaining rhodamine B dextran was determined from the stomach and segments of intestine to assess upper GI motility. SIT was estimated by the position of the geometric center of the rhodamine B dextran in the small bowel. The geometric center values are distributed between 1 (minimal motility) and 10 (maximal motility). n=10 in each group, representative data from three independent experiments are shown. **(i)** Whole gut transit time examined after five days of consecutive treatment of DT. An unabsorbable dye (carmine red) was administered by gavage and the time interval of first observance of the dye in stool was considered as whole gut transit time. n=10 in Piezo2-DTR and 6 in wild type control. Data are representative from two independent experiments. Error bars in panels e-i denote standard deviation of the mean; *p<0.05, ** p<0.01, *** p<0.001; unpaired two-tailed Student’s *t*-test.

In a 2-hour FPA, a 42% reduction in fecal pellet output was observed in the *Piezo2-*depleted mice compared to WT animals with the same treatment regimen (Fig. 6e). BEA demonstrated a substantial delay (36.9 ±19.1 min) to expel an inserted bead in the *Piezo2*-depleted mice compared to the WT controls (8.2 ±1.9 min) (Fig. 6f), which was not observed in the *Rosa26-LSL-DTR* animals under the same treatment (Supplementary Fig. 8g). Although gastric emptying (GE) was not affected in the *Piezo2-DTR* animals after DT treatment, small intestine transit (SIT) time, a measurement to assess the motility of small intestine, presented a small but statistically significant slowdown in the former group (Fig. 6g,h). There are a several possible explanations for this. Some Piezo2+ cells in the small intestine could have been depleted. Alternatively, 5-HT released from Piezo2+Tph1+ cells in the distal colon may provide feedback to the small intestine to accelerate motility, and thus depletion of these cells would result in slower intestinal transit. Consistent with the retarded colon motility, the whole gut transit time was found to be delayed in the *Piezo2-DTR* animals (181.8 ±17.6min) in comparison to WT (136.7 ±9.3 min, Fig. 6i) under the same DT treatment.

### Epithelial Piezo2 is important for normal colon motility

To directly test whether epithelial *Piezo2* is required to maintain normal colon motility, we used *Villin-cre* to deplete Piezo2 in gut epithelial cells. Unexpectedly, 15.9% of the *Villin-cre;Piezo2^fl/fl^*mice (referred to as Piezo2 CKO hereafter) died around 21-34 days after birth, affecting both males and females. By the time of humane euthanasia, the affected animals presented a 42% reduction of body weight and runt body size (Supplementary Fig. 9a-c).

*Piezo2* depletion was observed from the isolated epithelial layer of the distal colon, as assessed by qPCR (Fig. 7a) and smRNA-FISH analysis (Fig. 7b and Supplementary Fig. 9i). Anatomical and histological analysis suggested largely comparable intestine length and architecture of the gut wall with littermate controls (Supplementary Fig. 9d,e). Meanwhile, *Tph1* and *Chga* levels remained unaltered in the Piezo2 CKO animals assessed by qPCR (Fig. 7c and Supplementary Fig. 9f). Consistently, no change was observed in basal serotonin levels from either the epithelial tissue or the serum (Supplementary Fig. 9g,h). Notably, residual signals of *Piezo2* were observed in some of the *Tph1^+^* cells of the distal colon in the Piezo2 CKO animals (Fig. 7b), suggesting incomplete depletion. Importantly, *Piezo2* signals in the lamina propria remained largely unaltered (Fig. 7b and Supplementary Fig. 9i,j), suggesting that only the epithelial *Piezo2* was abolished in this mouse line.

**Figure 7.**
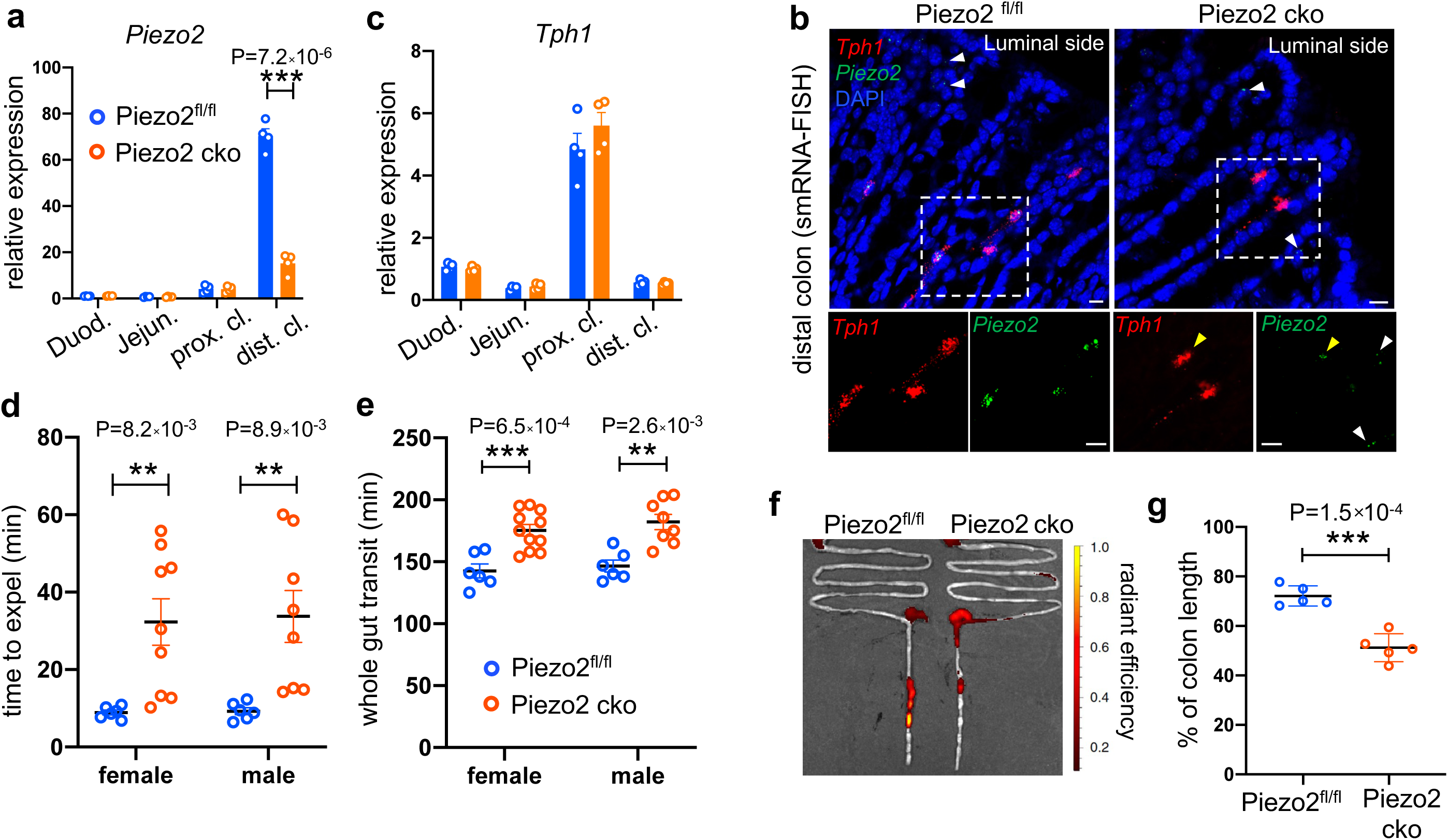
Epithelium *Piezo2* is required for efficient colon motility. **(a)** qPCR analysis of *Piezo2* depletion in *Villin-cre;Piezo2^fl/fl^* mice (Piezo2 cko) and *Piezo2^fl/fl^*. qPCR was performed on the RNA prepared from the epithelial extracts of the indicated regions of the gut. Gene expression was computed relative to the values in the *Piezo2^fl/fl^* duodenum, after normalization by the aggregates of three house-keeping genes (*B2m, Gapdh, Rpl13a*). Each circle represents one animal. Data were summarized from n=4 in each group. **(b)** smRNA-FISH assay of *Piezo2* and *Tph1* in either *Piezo2^fl/fl^* (left) or Piezo2 cko (right) mice. Dashed boxed are enlarged and presented in individual channels. Data are representative from five different animals per group. White arrowheads point to the submucosal signals of *Piezo2*. Yellow arrowheads point to the residual *Piezo2* signals in the *Tph1^+^* cells of the Piezo2 cko animals. Scale bars: 10 µm. **(c)** As in (a), qPCR analysis of *Tph1* in the Piezo2 cko and *Piezo2^fl/fl^* epithelium. **(d)** BEA. Each circle represents one animal. Representative data from two independent experiments. **(e)** Whole gut transit time. Each circle represents one animal. Representative data from two independent experiments. **(f)** Example fluorescent images of Piezo2 cko and *Piezo2^fl/fl^* intestine, 120 min after gavage of a fluorescent dye. **(g)** Summary data of fluorescent dye transit in the colon at 120 after gavage. % of colon length = dye travel distance in colon ÷ full length of colon × 100%. Each circle represents one animal. Representative data from two independent experiments. Error bars in panels a-e,g denote standard deviation of the mean; * p<0.05, ** p<0.01, *** p<0.001; unpaired two-tailed Student’s *t*-test.

**Figure 8.**
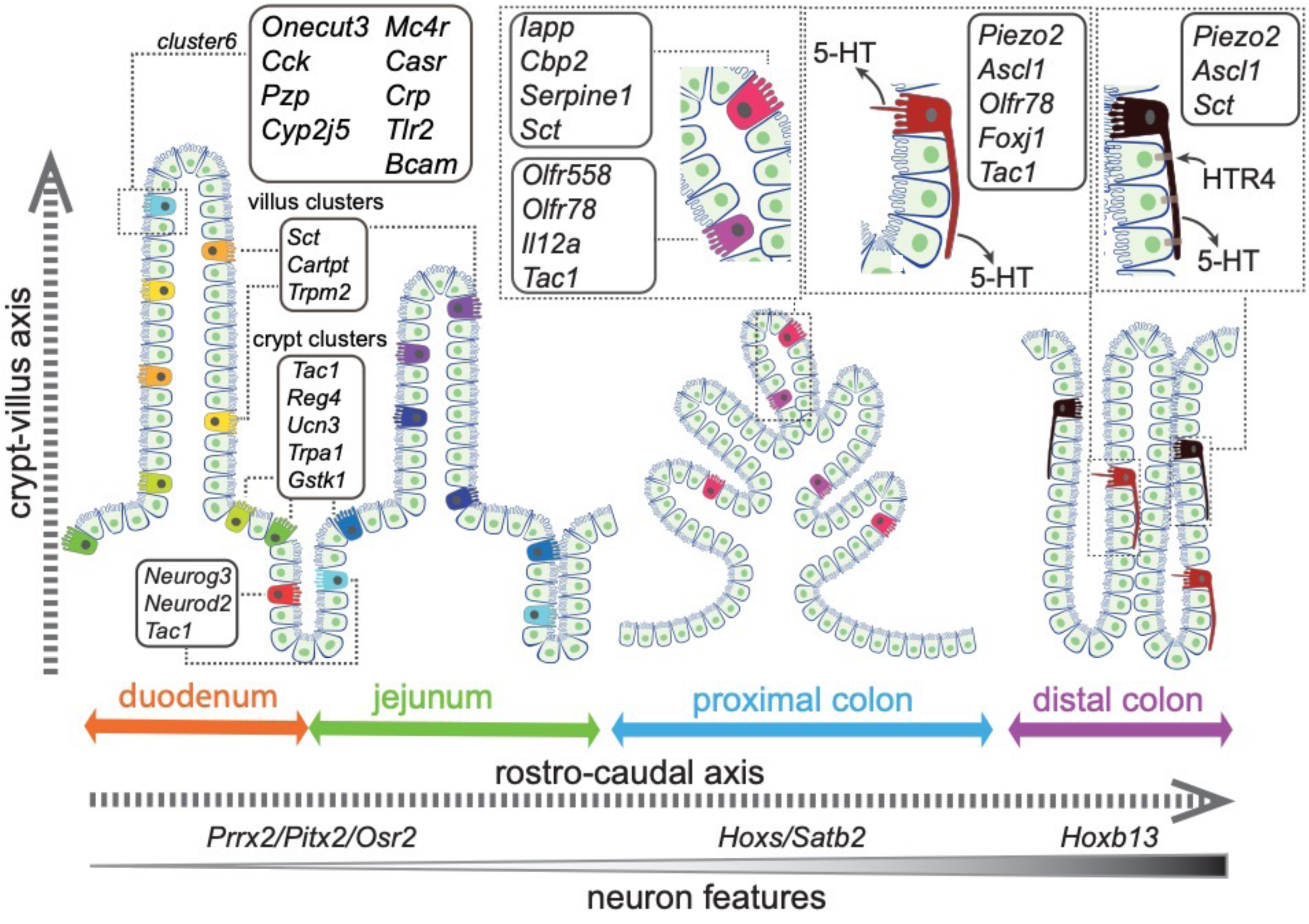
Summary of the spatial distribution of the identified EC clusters. Summary of the identified EC clusters and their respective gene signatures. Colors of the EC cells correspond to the clusters identified in this study.

Lastly, we measured BEA and total GI transit time. A significant slowing of expulsion was revealed by BEA in the Piezo2 CKO mice (male 33.7 ±19.0 min, female 32.3 ±18.1 min) compared with littermate Piezo2^fl/fl^ controls (male 9.25 ±2.2 min, female 8.9 ±1.5 min, Fig. 7d). In addition, prolonged whole gut transit time was observed in Piezo2 CKO mice (male 182 ±17.5 min, female 175 ±15.6 min) compared to littermate Piezo2^fl/fl^ controls (male 146 ±11.7 min, female 143 ±13.7 min; Fig. 7e). To assess small intestinal transit, mice were euthanized 70 min after gavage of a fluorescent dye and the travel distance of the dye within the intestine was calculated as a percentage of total small intestinal length. No difference was observed in small intestine transit between Piezo2 CKO mice (95.9 ±2.6%) versus Piezo2^fl/fl^ controls (96.6 ±2.2%, Supplementary Fig. 9k), in contrast to the DTR experiments, in which small intestinal transit was delayed. This could be due to the depletion of EC cells in the DTR experiments, whereas they are retained in the Villin-Cre Peizo2 KO mice. 5-HT secretion from ECs can be induced by other stimulants (even when Peizo2 is knocked out), and thus colonic 5-HT could be providing feedback to the small intestine to accelerate motility in the Villin-Cre Peizo2 KO mice. Residual Peizo2 expression in these mice could also be contributing to this effect. We then assessed fluorescent dye travel distance as a percentage of total colon length at 120 min after oral gavage, which was significantly shorter (51.2 ±5.6%) in the Piezo2 CKO mice compared to Piezo2^fl/fl^ controls (72.1 ±4.1%, Fig. 7f,g). Taken together, loss of Piezo2 in epithelial EC cells primarily affected colon motility.

Collectively, our *in vivo* gain- and loss-of-function analyses demonstrated that *Piezo2^+^/Ascl1^+^/Tph1^+^* cells are required for normal colon motility. By selectively targeting a subset of EC cells expressing specific sensors – here, a mechanical sensor – our study illustrates an example to effectively untangle pleiotropic functions of a complex cell population.

## DISCUSSION

In this study, we integrated high-throughput single-cell RNA-sequencing with spatial imaging analysis and constructed an ontological and topographical map for enterochromaffin (EC) cells in mouse intestine. We resolved 14 EC subpopulations characterized by their expression of distinct chemical and mechanical sensors, transcription factors, subcellular structures, and explicit spatial distribution within the gut mucosa (Fig. 7h). Together with *in vivo* functional validation of one subtype of EC cells, our study offers a framework to categorize complex sensory cells with defined molecular traits and led us to propose the functional identities for some of the subpopulations (Supplementary Table 1), while others warrant future investigation due to the limited information on their roles in GI biology (e.g. *Trmp2, Cartpt, Ucn3*).

We demonstrate that the transcriptome diversity of EC cells is closely related to their spatial distribution. The first layer of complexity is defined along the rostro-caudal axis, between the small intestine and the colon. Together with a host of TFs defining the rostro-caudal axis, we observe a gradual enrichment of a neuron-like transcriptome from small intestine to proximal and distal colon, such that *Ascl1*, encoding a key TF required for early neural fate commitment^49, 50^, is specifically expressed in the distal colonic EC cells. Concomitantly, these cells are featured with axon-like long basal processes and discrete expression of the mechanical sensor *Piezo2*. We resolved a second layer of complexity along the crypt-villus axis for the EC cells in a similar manner as described in EEC cells and enterocytes^23, 32^

By combining scRNA-seq profiling from two different genetic models, smRNA-FISH and *in vivo* genetic ablation, our study demonstrated that *Piezo2* is preferentially expressed in distal colonic EC cells that also have neuronal expression profiles. Neuron-like features are seen in the morphology of EC in the mouse distal colon, many of these having long processes, 50-100 µm or longer, that contain 5-HT^69^. Depletion of Piezo2 from these cells caused a 4-fold increase in bead expulsion time, implying that the mechanical stimulus provided by the bead caused 5-HT release and the initiation of colon propulsion. Consistent with this conclusion, in other studies in mice, 5-HT released from EC has been shown to cause colorectal propulsion^70^. In addition, *Piezo2* signals were observed in the lamina propria throughout the intestine. Given the previous report that Piezo2 was detected *via* scRNA-seq in a subset of sensory DRG neuron innervating the colon^71^, it is possible that both distal colonic EC cells and the sensory DRG neurons contribute to mechanosensory sensing and motility control. Such a dual component epithelial cell-neuronal sensory machinery parallels mechanisms described in the skin (Merkel cell-neurite complexes)^63, 72^, lung (neuroepithelial bodies)^62^ and most recently in the bladder (urothelial cells-sensory neurons)^64^. However, in contrast to Piezo2 signalling in ECs which results in accelerated gut transit, Piezo2 signalling in DRG neurons appears to slow transit^73, 74^.

A cluster of *Cck^+^/Oc3^+^/Tph1^+^* EC cells identified in the duodenal villi was enriched with two sets of sensory molecules: enzymes/receptors associated with pathogen/toxin recognition (*Crp*, *Lyzl4*, *Bcam*, *Tril*, *Tlr2*, and *Tlr5*) and receptors associated with nutrient sensing and homeostasis (*Casr* and *Mc4r*), suggesting a that these cells react to gastric content entering the duodenum. The gut has a well-established defensive role to expel noxious chemicals and toxins by nausea and vomiting that is initiated by 5-HT release and can be effectively inhibited by 5-HT3 receptor antagonists^75, 76^. Our findings suggests that *Cck^+^/Oc3^+^/Tph1^+^*cells equipped with pathogen/toxin recognition receptors may play a role in this defense mechanism. This includes C-reactive protein, encoded by *Crp* selectively expressed in *Cck^+^/Oc3^+^/Tph1^+^*cells, which is a conserved pattern recognition molecule involved in complement-mediated cell lysis^36, 77^ and lysozyme-like protein (LYZL) 4 that belongs to a family of antibacterial proteins, Toll-like receptors 2 and 5 and the Toll receptor interacting protein, TRIL. This suggests that the *Cck^+^/Oc3^+^/Tph1^+^* EC cells may react to pathogens both locally and through 5-HT signaling. Further study will be required to elucidate the molecular mechanism of this potentially important first line of defense. In regard to the nutrient-sensing molecules, previously, CasR and Mc4r have been reported in a subset of CCK^+^ I cells^78, 79^ and *Gcg^+^* L cells^80^, respectively, whereas *Pzp* is found to be expressed in a subset of *Gcg^+^* L cells^22^. Our integrated analysis indicates that all three molecules are enriched in the specialized *Cck^+^/Oc3^+^/Tph1^+^* cells.

EC cells reside along the frontier between the host and a highly dynamic range of chemicals and microorganism-derived signals within the intestinal lumen that are perturbed in various diseases, and, like other enteroendocrine cells, exhibit considerable plasticity^81, 82^. There are numerous reports of alterations to EC cell density in different pathophysiological states^75^ and, interestingly, some studies illustrating alterations to EC cell function. For example, EC cell sugar sensitivity is reduced in diet-induced obesity^83^, and colonic ECs in patients with ulcerative colitis have altered expression of genes relating to antigen processing and presentation and to chemical sensation^84^. Identification of orthologous EC subtypes in humans will be an important future step towards identifying how specific EC subtypes are affected in pathophysiological states, such as celiac disease, inflammatory bowel disease, and inflammatory bowel syndrome.

## METHODS

### Animals

All animal procedures performed at the University of California San Diego were conducted with approval by the Institutional Animal Care and Use Committee. All animal procedures at the University of Melbourne were conducted according to the National Health and Medical Research Council of Australia guidelines and were approved by the University of Melbourne Animal Experimentation Ethics Committee. *Neurod1-Cre* (Jackson Labs, 028364), *Rosa26-LSL-tdTomato* (Jackson Labs, 007914), *Piezo2-IRES-cre* (Jackson Labs, 027719), *Rosa26-LSL-CHRM3\** (Jackson Labs, 026220), *Rosa26-LSL-DTR* (Jackson Labs, 007900), *Piezo^fl/fl^* (Jackson Labs, 027720), *Vilin-cre* (Jackson labs, 021504) and C57BL/6 mice were purchased from Jackson Laboratories. Animals were housed in groups (2-5 mice/cage) in a specific pathogen-free facility provided with environmental enrichment (shelter, nesting material, *etc.*) and had normal immune status.

### Generation of *Tph1*-bacTRAP mice

A C57BL/6 BAC genomic clone RP23–4G4, which contains the locus of *Tph1* gene, was isolated from a RP23 mouse genomic BAC library (http://www.gensat.org/). BAC transgenic mice were produced according to published protocols^85^. The shuttle vector (S296-1) was digested with *Asc*I and *Not*I. A 420bp ‘A box’ fragment direct upstream of the ATG start codon of the *Tph1* gene was designed to be used for homologous recombination, amplified, digested with *Asc*I and *Not*I and cloned into the shuttle vector containing EGFP-RPL12. After electroporation, co-integration was identified by ampicillin resistance and verified by Southern blot using the A box sequence as the probe target. The modified BAC DNA was injected into fertilized oocytes of C57B6/J to generate the *Tph1*-bacTRAP line.

### Tissue dissociation and flow cytometry

Male mice aged 8-12 weeks were used. After the animals were sacrificed, small intestine and colon were surgically removed, rinsed with ice-cold PBS and the luminal contents flushed out with PBS using a 20 ml syringe with an 18-gauge round-tip feeding needle (Roboz Surgical Instrument, FN-7906). Duodenal and jejunal tissue was dissected between 1 to 5 cm and 13 to 17 cm distal of pyloric constriction. Ileum was dissected between 1 to 9 cm rostral of ileocecal junction. For the colon, the distal 4 cm tissue of descending colon was dissected as distal colon and the segment with banded lining distal to cecum was dissected as proximal colon. Dissected gut segments were inverted inside out and incubated in DMEM supplemented with 3 mM DTT (Sigma Life Science, D9779), 1 mM EDTA (Gibco, 15575-038) and 10% FBS (Gibco, 26410-079) at 37 °C for 30 min with consistent rotation. The released epithelial tissue was cut into smaller pieces, triturated with a 1000 µl pipette and dissociated in a collagenase solution with 1 U of Dispase (Stem Cell Technologies, 07923), 2 mg/ml Collagenase IV (Worthington Biochemical Corporation, LS004186) and 100 U DNase I (Worthington Biochemical Corporation, LS006330) in DMEM/F12 medium for 20-30 min at 37 °C with gentle mixing every 10 min. The dissociated cells were washed and filtered through a 100 µm cell strainer followed by a 40 µm cell strainer. The flow-through was spun down and filtered through another 40 µm cell strainer. The viability of the single-cell suspension was determined using trypan blue staining.

Cell pellets from the single-cell suspension were resuspended in FACS buffer (PBS, 5% FBS, and 5 mM EDTA) for staining in ice for 10 min with 7-AAD (BD Biosciences, #51-68981E). Only 7-AAD^−^ cells were considered as viable cells. To obtain cells for scRNA-seq, GFP^+^/7-AAD^−^ and GFP^−^/7-AAD^−^ cells were collected from the *Tph1*-bacTRAP mice, while tdTomato^+^/7-AAD^−^ and tdTomato^−^/7-AAD^−^ cells are collected from the *Neurod1-tdTomato* mice. Single-cell suspensions from different segments of gut were prepared and sorted separately. Single-cell suspensions from duodenum and jejunum were prepared from a single animal, while segments of proximal colon and distal colon were pooled from two and eight animals, respectively, to acquire sufficient numbers of cells. For ileum, however, even pooling eight animals did not yield adequate number for unbiased analysis (intestinal stem cells and transit amplifying cells were disproportionally enriched in the GFP^+^ cells, see data analysis), thus ileum was excluded from the subsequent scRNA-seq analysis.

### Library preparation and sequencing

Single-cell suspensions of freshly sorted cells were spun down to concentrate and were counted. All scRNA-seq libraries were prepared in parallel using Chromium Single Cell 3’Reagent Kits (10X Genomics; Pleasanton, CA, USA; *Tph1*-bacTRAP and small intestine of *Neurod1-tdTomato*: v2; colon of *Neurod1-tdTomato:* v3) according to the manufacturer’s instructions. Generated libraries were sequenced on an Illumina HiSeq4000 instrument, followed by de-multiplexing and mapping to the mouse genome (build mm10) using CellRanger (10X Genomics, version 2.1.1). Our sequencing saturation ranged between 61.0 and 81.7%.

### Multiplex fluorescent single molecule RNA *in situ* hybridization (smRNA-FISH)

To prepare tissue sections for RNA-FISH, wild-type C57BL/6 mice (8-10 weeks) were sacrificed, intestine and colon were dissected as described above, cleaned and fixed in 4% paraformaldehyde (PFA, Electron Microscopy Sciences, 15713) at 4 °C overnight, and cryoprotected in 30% sucrose (Ward’s Science, 57-50-1) for 24 h before being embedded in 100% O.C.T. Compound (Tissue-Tek). Cryosections were prepared at 12 µm. Single-molecule RNA-FISH was performed using the RNAscope. Multiplex Fluorescent Detection Kit v2 (323100, Advanced Cell Diagnostics) according to manufacturer’s instructions using TSA with Cy3, Cy5, and/or Fluorescein (Perkin Elmer NEL760001KT). All sections were counterstained with DAPI (1:1000; Invitrogen D1306). All co-staining was performed on tissue samples from at least three mice. Images were acquired on Zeiss LSM780 confocal microscope. Subsequent processing of images was performed in Fiji^86, 87^, including channel merging, pseudocoloring, maximum intensity projection, brightness adjustment.

### Quantification of smRNA-FISH

To quantitate crypt-villus distribution of cells, 9×9 tile images (1024-pixel × 1024-pixel) of an entire cross section of duodenum were acquired on Zeiss LSM780 and stitched in ZEN software. For automated analysis, images were preprocessed in Fiji (2.0.0) by maximum intensity projection, background subtraction and contrast enhancement, and the processed images were output as binary images for each channel. A custom CellProfiler pipeline was built to quantitate co-staining of two or three channels. Briefly, regions of interest (ROIs) were manually drawn on the DAPI channel in order to count cells separately in the crypts versus the villi. For each set of images, objects were identified independent in each channel, the signal of which was expanded by a 5-pixel diameter circle. If the circular objects identified from different channels overlapped, the cells were identified as double- or triple-positive. For each experiment, the pipeline was automated through images collected from at least 3 sections per animal in three animals. The number of double-positive cells (e.g., *Tph1+/Cartpt+*) was divided by single-positive cells (e.g., *Tph1^+^*) in each ROI and summarized across three animals to calculate mean and SEM statistics were performed using an unpaired Student’s *t*-test to compare enrichment in the crypts versus in the villi. To quantitate co-expression of two or three feature genes, the same pipeline was employed without demarcation of crypts versus villi.

### Quantification of Piezo2 depletion *via* smRNA-FISH

Sections were prepared from distal colon of either Piezo2-DTR or WT mice after intraluminal treatment of DT for 5 days. A custom CellProfiler pipeline was built to quantitate staining of *Tph1* and *Piezo2* channels. For each set of images, puncta (objects) were identified in the *Tph1* or *Piezo2* channel independently. When objects identified from the two channels overlapped, the puncta were identified as double positive; otherwise, the puncta were identified as single positive. The intensity of the *Piezo2* signals was quantitated by the size of puncta. Data were summarized from 4 pairs of animals with 2 tiled images from each animal. smRNA-FISH probes are listed in Supplementary Table 3.

### qPCR analysis

Cells were dissociated from the stripped epithelial layer in the duodenum, jejunum, proximal and distal colon of the *Tph1*-bacTRAP animals. Total RNA was prepared from the sorted regional GFP^−^ and GFP^+^ cells using the RNeasy Micro Kit (Qiagen). First-strand cDNA was synthetized from 100 ng RNA using the SuperScript III First-Strand Synthesis System (ThermoFisher, 18080051). Quantitative PCR was performed using FastStart Universal SYBR Green Master Mix (Sigma, 4913850001). The aggregates of three housekeeping genes (*B2m, Gapdh* and *Rpl13a*) were used to compute deltaCt. Relative gene expression was calculated by normalization against GFP^−^ cells extracted from the duodenum. Oligonucleotide sequences are listed in Supplementary Table 4.

### Tissue processing for immunohistochemistry

Duodenal (descending duodenum), jejunal (segment distal to root of mesentery), proximal colon (segment with banded lining distal to cecum), and distal colon (straight descending colon) tissue were isolated from three 2-month wild-type C57Bl/6 male mice and three 8-month old female *Tph1*-bacTRAP mice. The segments were opened along the mesenteric attachment, pinned onto balsa wood with the mucosal side facing up, and then fixed overnight in fixative (2% (v/v) formaldehyde plus 0.2% (v/v) picric acid in 0.1 M sodium phosphate buffer, pH 7.2) at 4 °C. Following fixation, tissue was washed three times in dimethyl sulfoxide, 10 min each, followed by three washes in PBS, 10 min each. Tissue was stored in PBS containing 0.1% (v/v) sodium azide until ready for use. Tissue was prepared for processing by placing segments in 50% (v/v) PBS-sucrose-azide and 50% (v/v) O.C.T. mixture for 24 h before being embedded in 100% O.C.T. Compound.

### Immunohistochemistry and image analysis

Sections (12 μm) were cut and allowed to dry at room temperature for 1 h on microscope slides (SuperFrostPlus; Menzel-Glaser; Thermo Fisher, Victoria, Australia). Sections were next incubated with 10% (v/v) normal horse serum prepared in PBS containing 1% (v/v) Triton-X-100 for 30 min, followed by overnight incubation at 4°C with primary antibodies (see Supplementary Table 2). Sections were washed three times in PBS and incubated with secondary antibodies (see Supplementary Table 2) for 1 h at room temperature. Following three washes with distilled water, sections were stained with Hoechst 33258 solution (10 µg/ml in distilled water) for 5 min to allow visualization of nuclei. Slides were washed with distilled water and coverslipped using non-fluorescent mounting medium (Dako, Carpinteria, CA, USA). Slides were allowed to dry overnight at room temperature after which they were imaged at 40× magnification using the AxioImager microscope (Zeiss, Sydney, Australia), or with the LSM800 (Zeiss) at 20× magnification. Immunoreactive cells were quantified by counting approximately 100 cells from each region of the gut for each of the 3 animals.

### Serotonin measurements

Blood samples were collected retro-orbitally from indicated animal cohorts. Animals subjected to blood collection were not used for *in vivo* motility assays. Serum was separated after coagulation at room temperature for 60 min, meaning that platelets could be contributing to the serotonin levels measured. Serotonin levels were detected in sera by ELISA according to the manufacturer’s instructions (Eagle Biosciences). To examine tissue serotonin levels from epithelial tissue, epithelium was extracted by incubation in DMEM supplemented with 3 mM DTT (Sigma Life Science, D9779), 1 mM EDTA (Gibco, 15575-038) and 10% FBS (Gibco, 26410-079) at 37 °C for 30 min with consistent rotation. Dissociated epithelial cells were lysed with standard buffer provided in ELISA kits. Cleared supernatant was used for ELISA. For the serotonin secretion assay, ∼1,000 sorted cells were equilibrated in standard buffer (+0.1% w/v ascorbic acid) for 30 minutes after wash with PBS. Cells were then incubated with standard buffer supplemented with indicated drugs or vehicle control, as well as 0.1% ascorbic acid for 15 minutes. Supernatant and cell lysates were prepared to measure secreted serotonin and cell lysate serotonin separately, the sum of which is calculated as total serotonin level. Secreted serotonin was considered as supernatant serotonin ÷ total serotonin × 100%.

### CNO and DT administration

Clozapine-N-oxide (Tocris, 4936) was dissolved in sterile 0.9% saline (Quality Biological, 114-055-101). On the day of experiment, the animals were anesthetized with isoflurane (5%, 1 l/min). CNO at indicated doses or saline control were administrated intracolonically using a 1 ml syringe with a 22-gauge round-tip feeding needle (round tip diameter 1.25mm, length 36mm; Roboz Surgical Instrument, FN-7920). Initially, two different doses (60 ng/kg, 120 ng/kg) of CNO were tested. As they were equally potent to induce serum serotonin elevation, the lesser dose (60 ng/kg in 50 µl) was chosen for the rest of experiments. Diphtheria toxin (DT; Sigma, D0564) was reconstituted in sterile water and administrated to anesthetized animals in the same manner as CNO, at 50 µg/kg in 50 µl twice a day for five consecutive days. Different regimens of DT administration were tested initially to obtain the maximal depletion of *Piezo2^+^* cells in the distal colon.

### *In vivo* motility assays

Animals were assigned to a random number before functional assays and data were collected in a blinded manner.

### Fecal pellet assay

The night before the assay, all animals (8-10 weeks, male) were housed singly in regular cages with wire mesh bottoms and bedding underneath with free access to food and water. After overnight acclimatization, each animal was placed into new wire mesh bottom cages. Fecal pellets were collected over 2 h and counted for each animal. For the cohorts receiving CNO treatment, since increased fecal pellets were already observed within the first 15 min after CNO administration in the *Piezo2-CHRM3\** mice, we have shortened the collection time window to 15 min for this experiment.

### Bead expulsion assay

All animals (8-10 weeks male or as indicated in the manuscript) were fasted for 2 hours before assay. Briefly, the animals were anesthetized with isoflurane (5%, 1l/min). A glass bead (2 mm in diameter, Sigma 1040140500) was placed into the colon using a disposable feeding needle with silicone tip (Fisherbrand, 01-208-89) to a distance of 2 cm from the anal verge. Mice were returned to their individual cages and allow to come back to full consciousness. Time required to expel the glass bead was monitored and recorded in each animal. The experiment was terminated at 60 min after mice became fully conscious. The mice failed to expel glass beads within this time window were reported as 60 min.

### Gastric emptying and small intestine transit analysis

All animals (8-10 weeks, male) were fasted overnight in cages that lacked bedding. Water was withdrawn 3 h before the experiment. Mice were orally gavaged with 100 µl sterile solution of 10 mg/ml rhodamine B dextran (Sigma R9379) in 2% methylcellulose (Sigma, M7027) through a 20-gauge round-tip feeding needle (Roboz Surgical Instrument, FN-7903). Animals were scarified 15 min after gavage; the stomach, small intestine, cecum, and colon were collected PBS. The small intestine was divided into 10 segments of equal length, and the colon (used to obtain total recovered rhodamine B fluorescence) was divided in half. Each piece of tissue was homogenized in PBS and centrifuged (2000 × *g*) to obtain a clear supernatant. Rhodamine fluorescence was measured in 250 µl aliquots of the supernatant (Tecan Trading, Infinite M200). Gastric empty rate was calculated as [(total recovered fluorescence − fluorescence remaining in the stomach) ÷ (total recovered fluorescence)] × 100%. Small intestinal transit was estimated by the position of the geometric center of the rhodamine B dextran in the small bowel^88^. For each segment of the small intestine (1–10), the geometric center (*a*) was calculated as follows: *a* = (fluorescence in each segment × number of the segment) ÷ (total fluorescence recovered in the small intestine). The total geometric center is Σ (*a* of each segment). Total geometric center values are distributed between 1 (minimal motility) and 10 (maximal motility).

### Total gastrointestinal transit time analysis

Male animals between 8-16 weeks were used, or as indicated in the manuscript when both female and male animals were examined. The night before the experiment, animals were transferred to individual housing with free access to water only. On the day of experiment, animals had free access to food and water for 1 hour. A solution of 6% carmine red (300 µl, Sigma, C1022) was prepared using 0.5% methylcellulose (Sigma, M7027) and was administered by gavage through a 21-gauge round-tip feeding needle (Roboz Surgical Instrument, FN-7903). 90 min after gavage, fecal pellets were monitored for the presence of carmine red. Total GI transit time was calculated from the time of administration to the first observance of carmine red in stool.

### In vivo transit analysis

Male animals between 8-16 weeks were used for this assay. Animals were fasted overnight and then had free access to food and water for 1 h. 100 µl Gastrosense 750 (Perkin Elmer) prepared in PBS was administered to the stomach via gavage. To assess partial intestinal transit, the GI tract (stomach to terminal colon) was removed at indicated time. Fluorescence images were obtained of the GI tract using an IVIS Lumina II In Vivo Imaging System (Perkin Elmer).

### scRNA-seq data processing

#### Gene expression matrices

Gene expression matrices were generated using the CellRanger software (10X Genomics). Sample data were aggregated using CellRanger (cellranger -aggr) and resulting data were processed further in Python (version 3.6.2) or scanpy (version 2.1.4)^89^.

#### Quality control

The following quality control steps were performed: (1) Non-coding gene and genes expressed in fewer than 10 cells were not considered; (2) cells that expressed fewer than 500 genes were excluded from further analysis (suggesting a low quality of cells); (3) cells in which >10% of unique molecular identifiers (UMIs) were derived from the mitochondrial genome were removed.

#### Cell doublet removal

Since we kept our capture rate low (∼2,000 cells per sample) during library preparation, the cell doublet rate was low as expected. We removed potential cell doublets based on: (1) the presence of gene signatures from two different cell classes, such as epithelial markers and immune cell markers; (2) the observation of a second peak of total UMIs distribution in comparison to the cells from the same class.

#### Normalization

Given the presence of stochastic zeros in scRNA-seq data and the wide distribution of total UMIs per cells in our dataset, we used a pool-based size factor to deconvolute cell-specific size factor for each single cell library for normalization purpose. Briefly, a coarse cell clustering was first performed using hierarchical clustering based on the top 50 components of principle component analysis (PCA) for the entire expression matrix. Summation of UMI counts across cells in each resolving cluster was computed as pooled-based size factor and repeated to generate a linear system for all single cells. A weighted least-squares approach is applied to solve the linear system and to deconvolute a cell-base size factor for all single cells, as implemented in Scran (version 2.1.6)^90^. The UMI counts were then normalized by a cell-specific size factor and transformed as log_2_(normalized counts +1). For simplicity, the transformed normalized counts are presented as log_2_(counts).

#### Clustering and spatial visualization

Linear dimensionality reduction was performed on the aggregated dataset using principal component analysis (PCA). The top PCs were chosen based on elbow plots, where the percentage variance explained by each PC was plotted, and the number of principal components was chosen as a substantial drop was observed in the proportion of variance explained. Typically, 20-30 PCs were chosen based on our dataset, and were visualized using t-distributed Stochastic Neighbor Embedding (t-SNE)^91^. Graph-based clustering was performed for community detection. Briefly, a k-nearest neighbors (kNN) graph was built based on the Euclidian distance of the single cells in the PCA space, the edges between the detected community were weighted using Jaccard similarity, and the Louvain method was applied to optimize the modularity of the communities. Cluster numbers were chosen based on Bayesian information criterion (BIC) and biological considerations. Cluster resolution (parameter of k), t-SNE perplexity and the number of PCs used for clustering and visualization were adjusted based on the total cell number of the dataset. (Related to Fig. 1d,h,g; Supplementary Fig. 1c,d,e,g; and Fig 3i).

#### Identification of EC cells and non-EC cells

In the initial scRNA-seq profiling from the *Tph1*-bacTRAP mice, we obtained 4,729 signal cells including GFP^−^ (2,412) and GFP^+^ (2,317) cells. Clustering analysis coupled with spatial visualization in t-SNE space, as described above, indicated ∼23% of the GFP^+^ cells clustered together with the GFP^−^ cells. To annotate GFP^−^ cells and GFP^+^ clustered with GFP^−^ cells, we obtained gene sets from GSE92332^20^, which including gene sets associated with each major cell types in intestinal epithelium have been identified, including: intestinal stem cells, transit amplifying cells (TACs), immature enterocytes, mature enterocytes, tuft cells, goblet cells and enteroendocrine cells. We calculated module scores for each cell cluster identified in our dataset by computing the average expression levels of each cell type gene set subtracted by the aggregated expression of all detected genes in our dataset. Cell types were assigned based on the highest module scores across all cell types as described above. Furthermore, we applied z-score transformation to the cell type-enriched gene set and validate the cell type assignment based on their respective z-score enrichment. The same approaches were employed to identify the non-EC cell types in the second scRNA-seq profiling from the *Tph1*-bacTRAP mice, thus annotated intestinal stem cells, TACs, immature enterocytes, mature enterocytes and colonocytes. The T lymphocytes and mast cells were identified based on the top differentially expressed genes in the respective clusters. (Related to Fig. 1d and Supplementary Fig. 1d).

#### Elimination of ileal GFP^+^ cells

Previous studies have indicated a significant decline of 5-HT^+^ cells in the distal small intestine^92^. Consistent with such observation, in the initial scRNA-seq profiling of the single epithelial cells from *Tph1*-bacTRAP mice, we obtained only 225 GFP^+^ single cell libraries from the ileal epithelial cells collected from eight *Tph1*-bacTRAP mice, which was significantly lower than those from other regions of the gut (0.1% GFP positivity compared to ∼0.3-0.5% positivity in other regions). Additionally, the total number of epithelial cells obtained from the ileum in each mouse was only 20% of those from duodenum, contributing to the lower number of GFP^+^ cells obtained from the ileum. Upon further computational analysis, as much as 72% (162) of these GFP^+^ cells clustered with non-EC cells, in particular, 49% (111) were identified to have marker genes for stem cells or TACs, indicating that a majority of these cells are not fully committed EC cells. We thus eliminated ileum from further analysis.

#### Identification of differentially expressed genes in cell populations

To identify genes expressed at significantly higher level in one cluster or region than the other clusters or regions, we used the Wilcoxon rank-sum test, which is non-parametric and does not assume normality. Correction for multiple testing was performed using the Benjamini-Hochberg procedure to control the false discovery rate (FDR). We ran differential expression tests between each pair of clusters (regions) for all possible pairwise comparisons, as implemented in Scran (version 2.1.6). For a given cluster (region), the DE genes were filtered using the maximum FDR q-values across all pairwise comparisons. Genes that are known to be associated with dissociation process were not considered to be to be differentially expressed genes^93^. For Fig. 1l, 2f, 4e, DE genes were obtained using maximum FDR<10^−10^.

#### Identification of signature genes for clusters

To identify maximally specific genes for each EC cell clusters, we performed pairwise differential gene expression analysis as described above between each pair of clusters for all possible pairwise comparisons. For a given cluster, putative signature genes were ordered based on FDR (FDR<10^−10^, the smallest FDR was ranked on top). The final signature genes lists were obtained by calculating the rank product for selected genes in all pairwise comparisons. Rank product statistic^94^ was used to determine p values for each marker gene and was adjusted by the Benjamini-Hochberg procedure to correct multiple hypothesis testing. To ensure the signature genes are enriched in EC cells but not other cell types in gut epithelium, the candidate genes were evaluated against both GFP^−^ cells and non-EC cells identified in our own datasets and against the GSE92332 dataset^20^.

#### GO term enrichment analysis

Differentially expressed genes identified by regions (Fig. 1h) or by clusters (Fig. 2h and Supplementary Fig. 2i) were selected by FDR<10^−10^ and subjected to enrichment analysis by accumulative hypergeometric test followed by Sidak-Bonferroni correction as implemented in Metascape^95^. The resulting GO terms were selected by q-values that are presented by the size of the hexagons. The number of DE genes identified in the GO terms are represented by heatmap.

#### Comparison with public datasets

We compared neuropeptidergic hypothalamic neurons with enterochromaffin (EC) cells based on two considerations: 1) hypothalamic neurons produce many hormone peptides similar to gut enteroendocrine cells, and 2) colonic EC cells are enriched with many neuronal signatures. We extracted expression matrices and metadata from GSE74672^53^. Cell types were catalogued as indicated by metadata. Data were processed in the same pipeline as described for the *Tph1*-bacTRAP or *Neurod1^+^* dataset. DE gene analysis was performed to identify neuron-enriched genes against both microglia and oligodendrocytes (FDR<10^−10^). The resulting gene set was used to compare EC cells originated from different segments of the gut. To identify orthogonal signature genes from human gut mucosa, we cross-compared the *Neurod1^+^*dataset with GSE125970^21^, where human gut mucosal cells were isolated from biopsy samples in ileum, colon and rectum. Since no selection has been applied to dissociated human epithelial cells, <1% of captured cells are enteroendocrine cells. We identified 63 L cells and 34 EC cells in total, which were determined as *Pyy^+^, Gcg^+^* or *Pyy^+^/Gcg^+^* for the former, and *Tph1+* for the latter. Data were processed in the same manner as *Tph1*-bacTRAP or *Neurod1^+^* dataset.

### Code and data availability

Data used to generate figures and graphs are available within the article and in the Source Data files. Scripts used generate the figures will be made available from the corresponding authors upon reasonable request. Raw fastq files and Cell Ranger processed digital gene expression matrixes (DGE) have been deposited at NIH’s Sequence Read Archive (SRA) under SUB7225910.

### Statistics

In addition to the statistic tests described in the data analysis section, the following tests were performed:

For Fig. 1I, Fig. 2j and Supplementary Fig. 2g,h: A two-sample Kolmogorov-Smirnov test was performed to test whether two underlying probability distributions are the same. Implemented by scipy.stats.ks_2samp in python.

For Fig. 2d: An unpaired two-tailed Student’s *t*-test was employed to test whether the positive fractions identified in the villi are significantly different from the ones observed in the crypts.

For Fig. 2i: Hypergeometric testing for enrichment of indicated hormones in the *Cck^+^*/*Tph1^+^*population versus the Tph1^+^ population was performed and implemented by scipy.stats.hypergeom in Python.

## ACKNOWLEDGEMENTS

We thank Ardem Patapoutian for conducive discussions. We acknowledge the help of Jesus Olvera and Cody Fine with FACS and Elsa Molina for assistance with confocal microscopy. This work is partially funded by a grant from the Takeda-Sanford Innovation Alliance. Sequencing was conducted at the Institute for Genomic Medicine (IGM) Genomics Core at UC San Diego which is supported by P30CA023100. M.P. was supported by National Research Service Award (NRSA) grant F32HL143978.

## AUTHOR CONTRIBUTIONS

G.W.Y, J.W. and Y.S. conceptualized the study. Y.S., and K.S.L. performed EC isolations and 10X Genomics single-cell sequencing. Y.S. processed scRNA-seq data and performed computational analysis. Y.S., K.S.L and B.L. performed smRNA-FISH experiments, data acquisition and analysis. Y.S. and M.P. performed animal experiments. A.K., L.J.F., S.D. and B.C. performed and quantified immunohistochemical localizations. J.H. shared reagents. Y.S., L.J.F., J.B.F., A.D. J.W. and G.W.Y. wrote the manuscript. All authors discussed and commented on the manuscript.

## FIGURES AND FIGURE LEGENDS

**Supplementary Figure 1.**
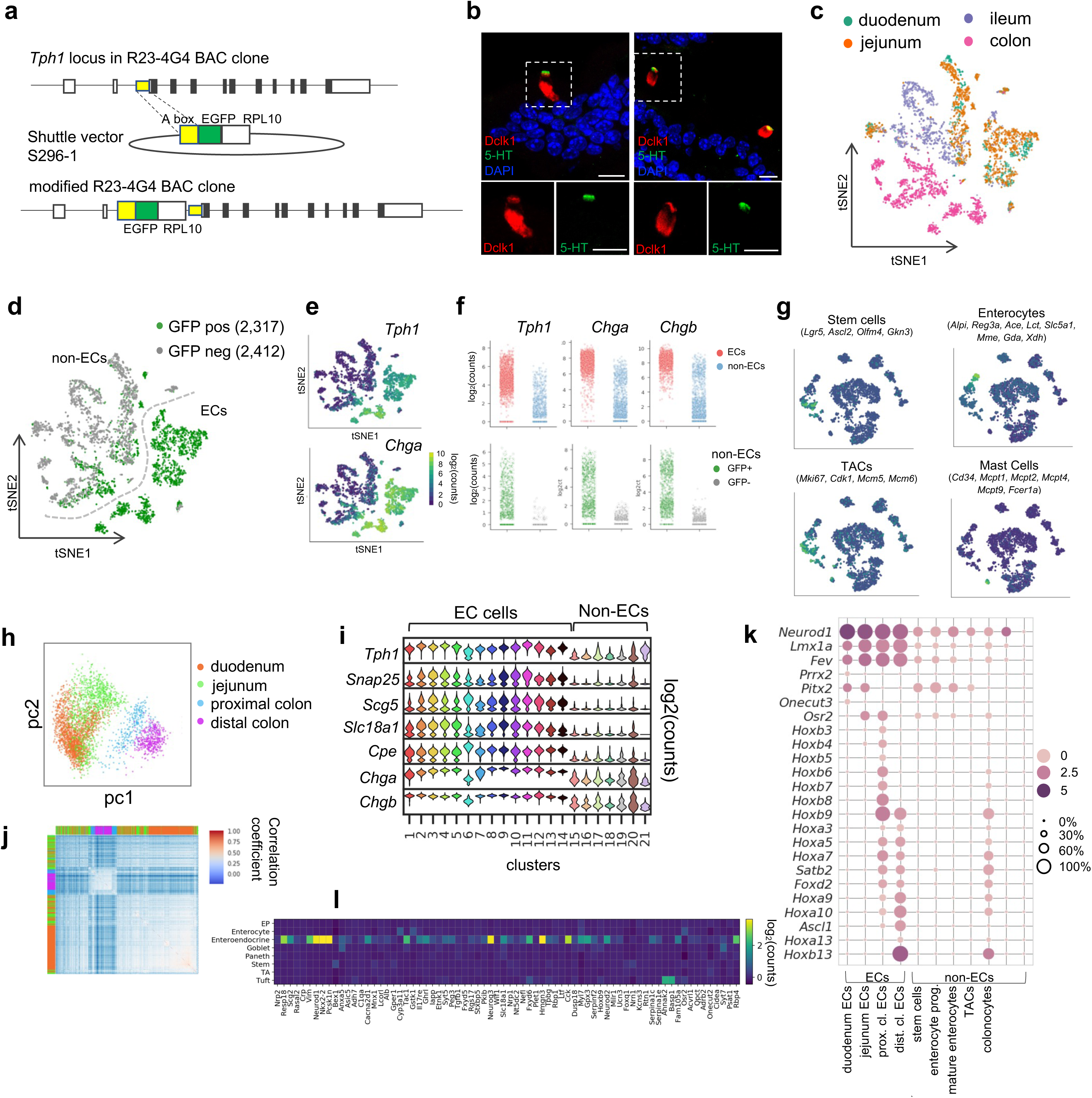
scRNA-seq identifies distinct intestinal enterochromaffin (EC) cell clusters. **(a)** Schematic of *Tph1-bacTRAP* allele generation. A R23-4G4 BAC clone containing the *Tph1* locus was used. Yellow box: ‘A box’, a 420 bp fragment immediately upstream of start codon of *Tph1* that was used for homologous recombination. A sequence encoding a fusion protein of EGFP and RPL10 was cloned immediately downstream of the A box. **(b)** Dual IF staining of 5-HT and Dclk1 (tuft cell marker) in the duodenum of wild type mice. Scale bar: 10 µm. **(c,d)** t-SNE projections of all cells (4,729) from single-cell profiling capturing both GFP^+^ (2,317) and GFP^−^ (2,412) cells from the indicated regions of *Tph1*-bacTRAP mice. Single cells are color-coded by regions in (c) or by GFP positivity in (d). **(e)** t-SNE projections of cells shown in (c,d) superimposed on heatmaps representing normalized counts for the indicated genes. **(f)** Comparison of *Tph1*, *Chga* and *Chgb* expression in EC cells versus non-EC cells (*upper*) or in the GFP^+^ versus GFP^−^ cells among non-EC cells (*lower*). **(g)** t-SNE projection of cells from the second profiling capturing GFP^+^ cells from the duodenum, jejunum, proximal and distal colon as presented in Fig. 1d,g,h. Superimposed heatmaps represent the aggregated transcript counts of the indicated marker genes for each identified cell type. TACs: transit amplifying cells. **(h)** PCA plot of EC cells (as shown in Fig 1d,g,h) from the second profiling and color-coded by regions. **(i)** Violin plots showing expression of well-established marker genes of EC cells in all clusters. **(j)** Unsupervised hierarchical clustering of EC cells based on transcriptome similarity (correlation coefficient) computed from the top 800 genes extracted from PC1 and PC2 (as shown in h). Color bars indicate the regions of EC cells. **(k)** Transcription factors (TFs) differentially detected along the rostro-caudal axis are presented in all cells (as shown in Fig 1d,g,h) partitioned by cell types or by regions (for ECs). Size of the circles represents percentage of expression and intensity of the circles represents aggregated expression of indicated TFs **(l)** Signature genes of the EC clusters (as shown in Fig 1i) showing enrichment in EEC cells but not in the other epithelial cell types of the intestine. Heatmap illustrates median expression of the indicated genes extrapolated from GEO dataset GSE92332. EP: early progenitor, TA: transit amplifying cell.

**Supplementary Figure 2.**
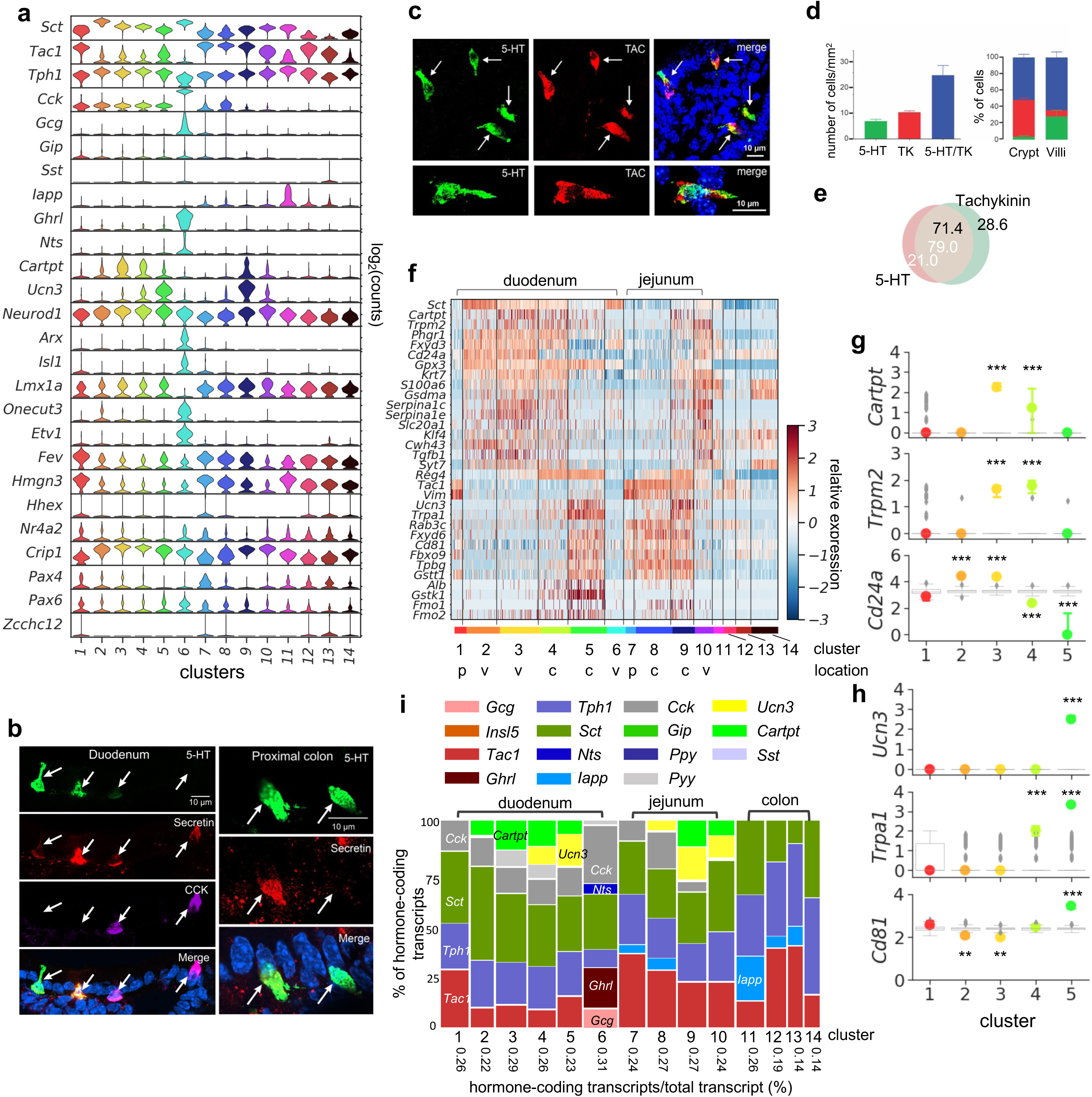
SI EC cells are predicted to switch sensors and hormone compositions along the crypt-villus axis. **(a)** Expression (log_2_(counts)) of peptide hormones along with *Tph1* and transcription factors in the EC clusters (as shown in Fig. 2a). **(b)** Co-expression of secretin, 5-HT and CCK examined by IF staining. **(c,d)** Co-expression of tachykinin and 5-HT examined by IF staining (c) and quantitated (d) in the jejunum **(e)** Venn diagram showing co-expression of tachykinin and 5-HT in the jejunum based on IF staining. Numbers present the percentage out of all counted 5-HT^+^ cells (white) or tachykinin^+^ cells (black). **(f)** Heatmap of genes exhibiting differential enrichment along the crypt-villus axis. Relative expression (z-score) of indicated genes are shown across all single EC cells. Color-coded bar at the bottom represents the clusters. p: progenitor, v: villus, c: crypt. **(g,h)** Representative gene enrichment in villus clusters (g) or crypt clusters (h). Note that cluster 4 shares some features in common with crypt and villus clusters and may represent cells at an intermediate stage of development. Point plots depict the median expression of indicated genes in each cluster and the error bars represent upper and lower quantiles. Clusters are represented by colors. The grey box plots depict the distributions of the median expression based on size-matched randomization control experiments (median of n=500 randomizations). The boxes present the quantiles and whiskers show the rest of the distribution. *** p<10^−50^, ** p<10^−30^; two-tailed Kolmogorov-Smirnov statistic between the observed gene distribution and randomized control distribution. **(i)** Marimekko chart showing the cluster-wise distribution of hormone-coding transcripts (color-coded) in the 14 clusters of EC cells. The width of each column represents the percentage, stated at the bottom, of all hormone-coding transcripts (plus *Tph1*) out of all transcripts. Height of each bar represents the percentage of indicated hormone-coding transcripts out of all hormone-coding transcripts.

**Supplementary Figure 3.**
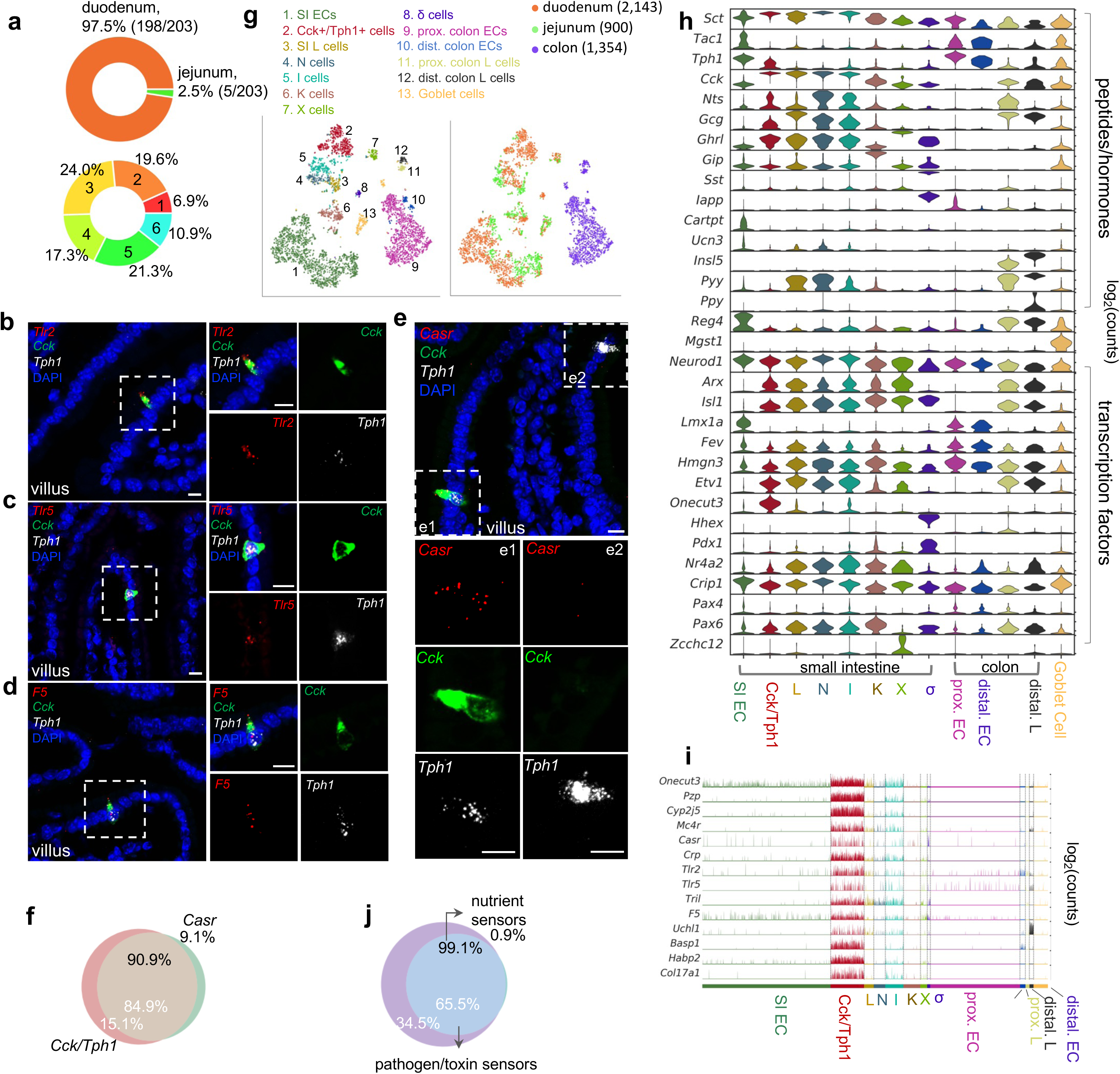
*Cck*, *Oc3* and *Tph1* specify an EC subpopulation with a dual sensory signature. **(a)** *Upper*: Regional composition of cluster 6 cells. Regions are color-coded. 203 cells were identified in cluster 6 in the *Tph1-bacTRAP* dataset. *Lower*: EC cells in duodenum are constituted by 6 clusters. Numbers and colors indicate clusters. **(b-e)** smRNA-FISH of *Tlr2*, *Cck*, and *Tph1* (b); *Tlr5*, *Cck*, and *Tph1* (c); *F5*, *Cck*, and *Tph1* (d); and *Casr*, *Cck*, and *Tph1* (e). Two dashed boxes (e1 and e2) show a representative *Cck^+^/Tph1^+^* double-positive cell stained positive for *Casr* (e1) and a*Tph1^+^* single-positive cell without *Casr* (e2). Scale bars: 10 µm. **(f)** Quantitation of *Casr* (black text) and double *Cck^+^/Tph1^+^*positive cells (white text) in the duodenum based on smRNA-FISH. 64 cells were quantitated from four mice. **(g)** tSNE representation of single *Neurod1-tdTomato^+^* cells isolated from the gut. Cells are color-coded by clusters identified *via* Louvain method (*left*) or by regions (*right*). **(h)** Expression (log_2_(counts)) of peptide hormones along with *Tph1* and transcription factors in the EEC clusters. **(i)** Expression of signature genes identified in the *Cck^+^/Tph1^+^* cluster from the *Neurod1-tdTomato* dataset. Cells are shown in the columns and color-coded by clusters, and genes are shown in the rows. **(j)** Venn diagram showing the co-expression of two sets of genes summarized by the aggregated expression of genes associated with pathogen/toxin recognition (*Crp*, *Lyz4*, *Tril*, *Tlr2*, and *Tlr5*) versus genes associated with nutrient sensing and homeostasis (*Mc4r* and *Casr*) in the *Neurod1-tdTomato* dataset.

**Supplementary Figure 4.**
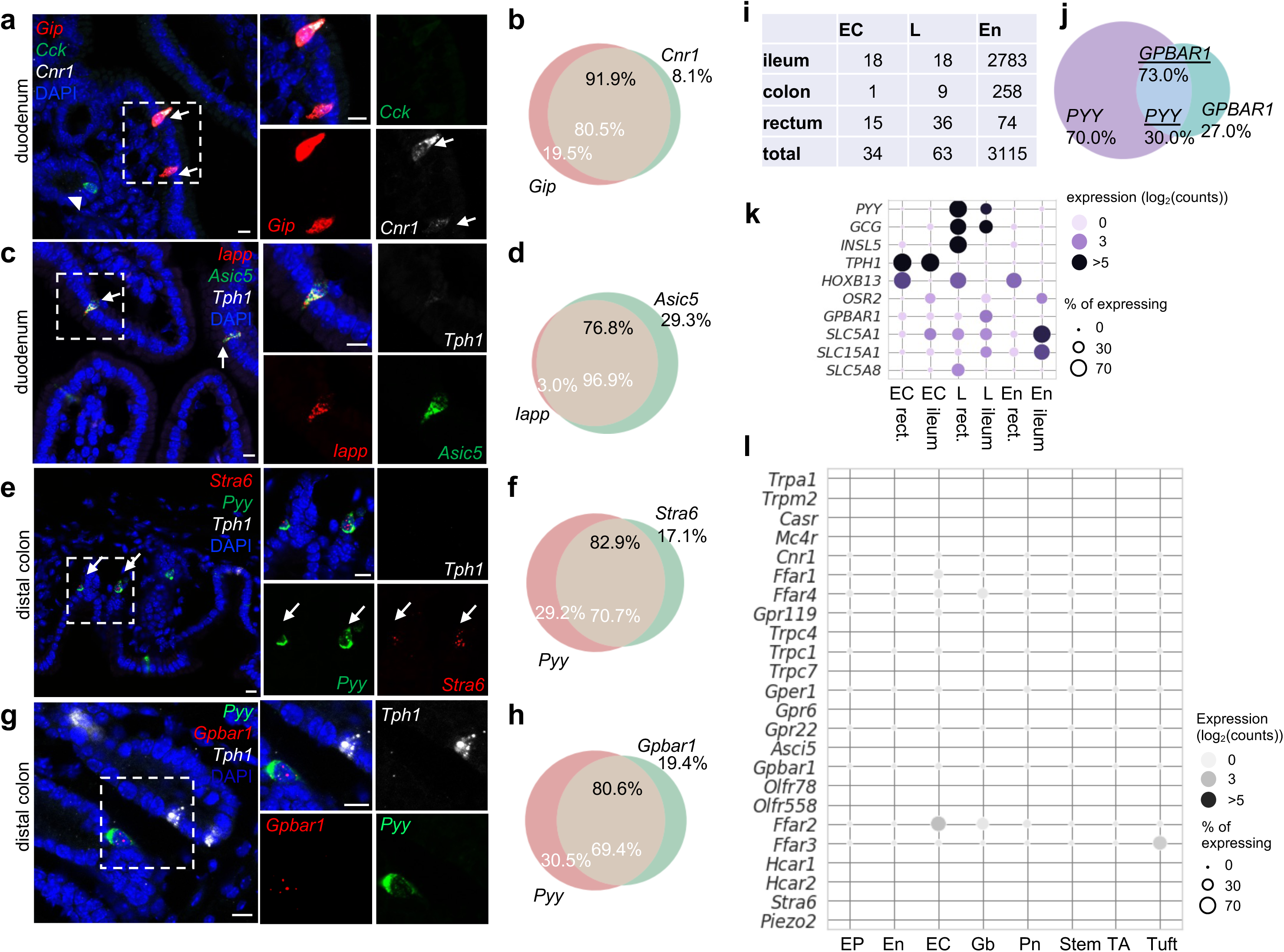
Distinct molecular sensors identified in EC cells versus other EEC cells. **(a-h)** smRNA-FISH of *Cnr*, *Gip*, and *Cck* (a); *Asic5*, *Iapp*, and *Tph1;* (c) *Stra6*, *Pyy*, and*Tph1* (e); or *Gpbar1*, *Pyy*, and *Tph1* (g) in wild type mice. Venn diagrams of the corresponding quantitation based on smRNA-FISH experiment (b, d, f, h). For each quantitation, 105-156 cells were counted from three mice. Scale bars: 10 µm. **(i)** Breakdown of single human mucosal cells (GEO dataset GSE125970) in indicated categories. En: enterocytes. EC: enterochromaffin cells. L: *Gcg* and/or *Pyy* positive L cells. Since only 10 EEC cells were identified in the colon, they were not assessed in the subsequent analysis. **(j)** Venn diagram of *GPBAR1* and *PYY* co-expression in human gut mucosal cells. Data were extracted from GEO dataset GSE125970. **(k)** Dot plot of key sensors differentially expressed in human ileal versus colonic L cells. Data were extracted from GSE125970. Single cells that are positive for *TPH1*, or positive for *PYY* or/and *GCG* are annotated as EC or L cells respectively. En: enterocytes. Enterocytes and regions are annotated as in the provided metadata. Note, *HOXB13* or *OSR2* expression mark rectum or ileum origin respectively. **(l)** Dot plot showing the expression level and percentage of sensors identified in EECs (Fig. 3j). Data were extrapolated from GSE92332. Note, many of sensors (10/24) identified in our study were not detected in GSE92332, possibly due to the small number of EEC cells (258) in this dataset. EP: early progenitor, En: Enterocyte, EC: Enterochromaffin cell, Gb: goblet cell, Pn: Paneth cell, TA: transit amplifying cell.

**Supplementary Figure 5.**
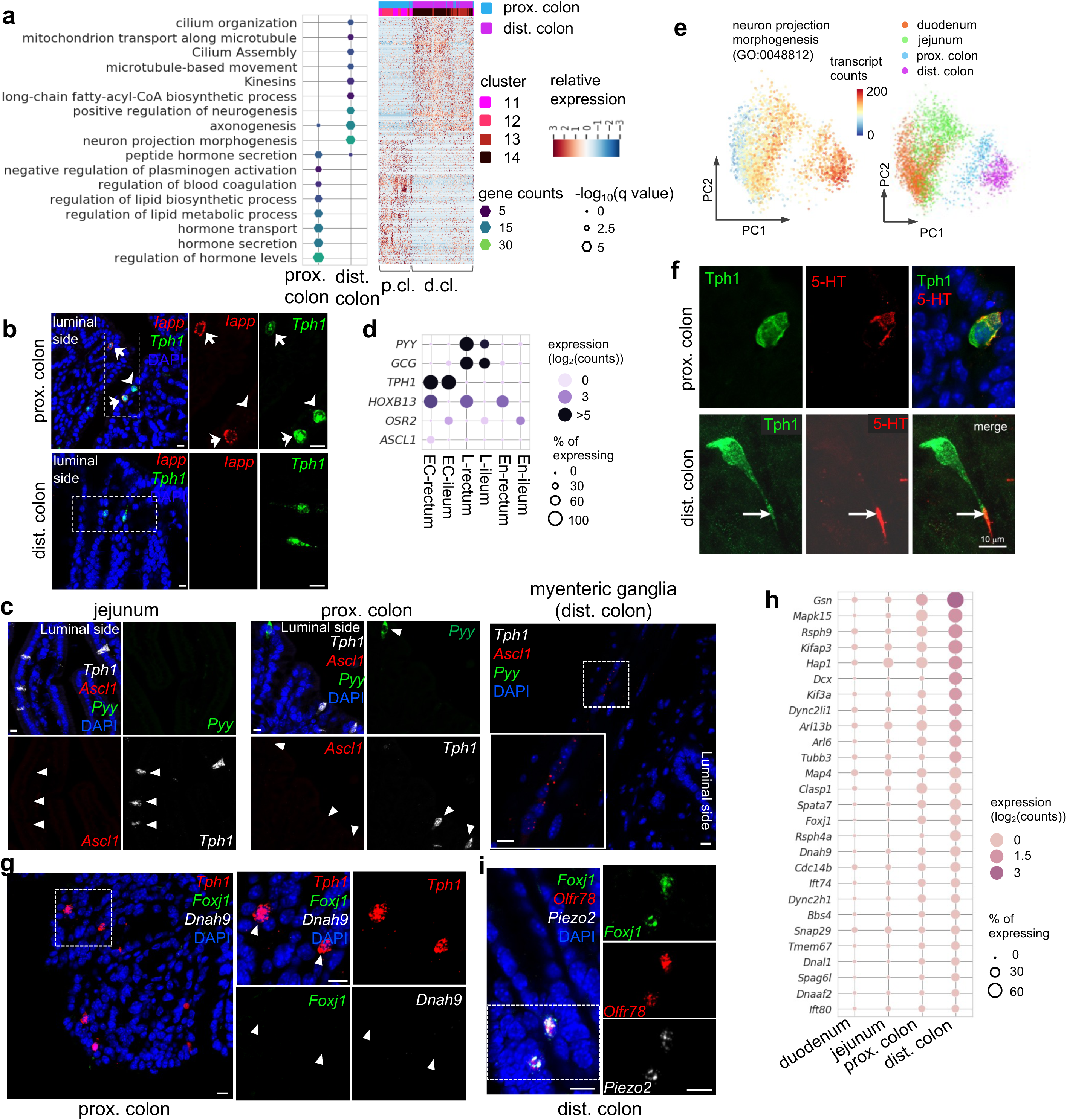
Distinct subpopulations of EC cells are resolved in the colon. **(a)** GO analysis of DE genes identified between the proximal and distal colonic EC cells (*left*). Data are based on the *Tph1-bacTRAP* dataset. DE genes: FDR<10^−10^, Wilcoxon rank sum test corrected with the Benjamini-Hochberg procedure for multiple hypothesis testing. Unsupervised hierarchical clustering of single cells (in columns) based on identified DE genes is shown on the *right*. **(b)** smRNA-FISH of *Iapp* and *Tph1* in the proximal colon (upper) and distal colon (lower). Arrows point to *Iapp/Tph1* double-positive cells. Arrowheads point to the absence of *Iapp* in some *Tph1^+^* cells. **(c)** smRNA-FISH of *Ascl1*, *Tph1*, and *Pyy* in the jejunum, proximal colon and myenteric ganglia from distal colon. Arrowheads point to the absence of *Ascl1* in either *Tph1^+^* or *Pyy^+^* cells. Associated with Fig. 4d. **(d)** Dot plot showing *ASCL1* enrichment in human rectal EC cells. Data were extracted from GEO dataset GSE125970. Single cells that are positive for *TPH1*, or positive for *PYY* and/or *GCG* are annotated as EC or L cells respectively. En: enterocytes. Enterocytes and regions are annotated according to the provided metadata. *HOXB13* and *OSR2* expression mark the origins of rectum and ileum respectively. Size of the circles represents percentage of expression and intensity of the circles represents aggregated expression of indicated genes. **(e)** PCA plots color-coded by the aggregated transcript counts of the genes (based on the *Tph1-bacTRAP* dataset) in the indicated GO term (*right*) and by regions (*left*). **(f)** IF staining against 5-HT and GFP (representing Tph1) in the proximal colon (upper) and the distal colon (lower). Note the axon-like extension is not observed in the proximal colonic EC cells. **(g)** smRNA-FISH of *Tph1*, *Foxj1*, and *Dnah9* in the proximal colon. The boxed region is enlarged on the right. Arrowheads point to the absence of *Foxj1 and Dnah9* in *Tph1^+^*cells in the proximal colon. Associated with Fig. 4h. **(h)** Dot plot showing expression levels and frequency of genes associated with ‘cilium assembly’ (R-MMU-5617833) and ‘cilium organization’ (GO:0044782) in EC cells along the rostro-caudal axis of the gut. **(i)** smRNA-FISH of *Foxj1*, *Olfr78*, and *Piezo2* in the distal colon. The boxed region is enlarged on the right and split into individual channels. Scale bars in panels b,c,f,g,i: 10 µm. Images are representative from three mice.

**Supplementary Figure 6.**
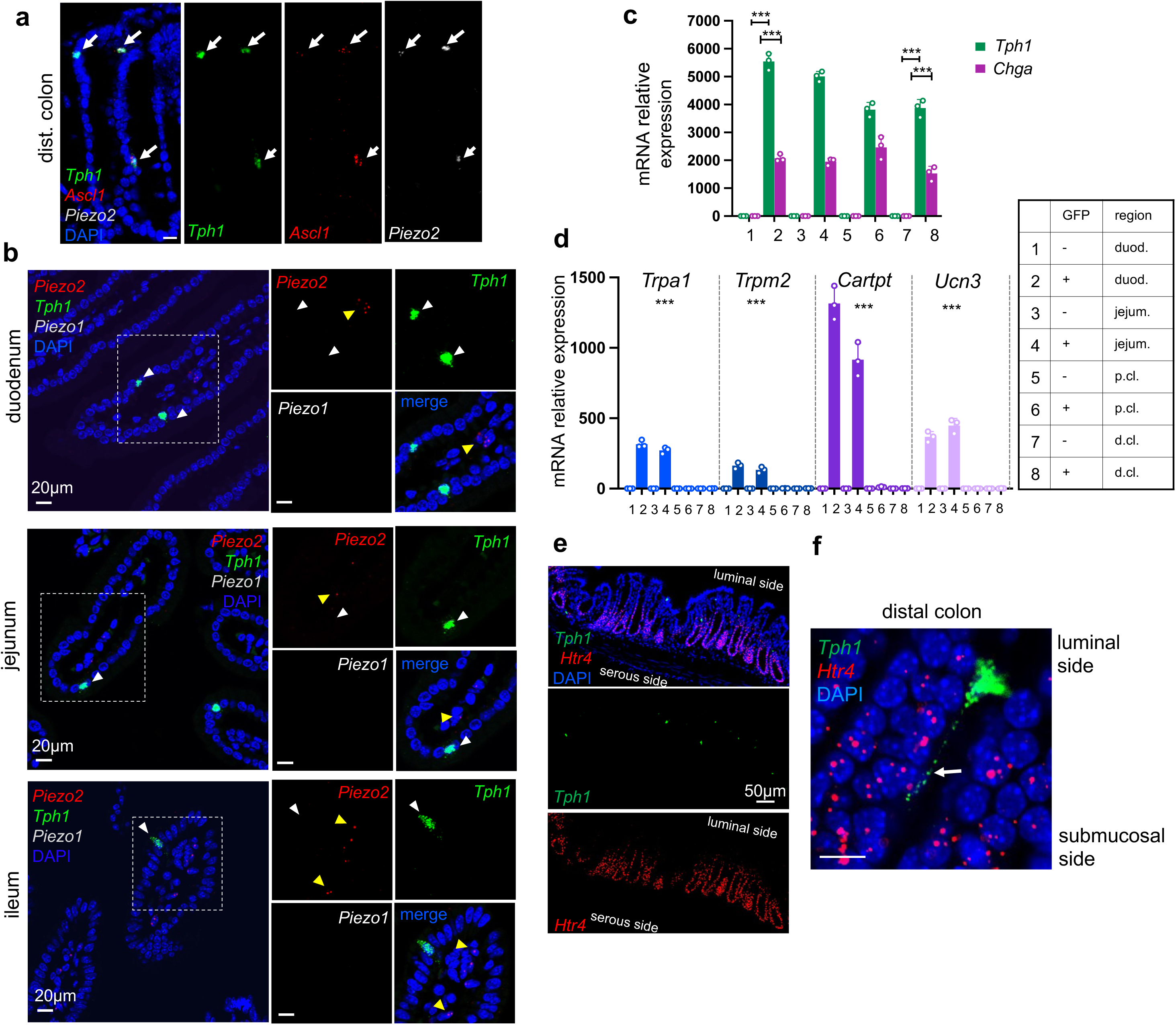
*Piezo2* is highly enriched in distal colonic EC cells that mediate colon motility. **(a)** smRNA-FISH of *Tph1*, *Ascl1*, and *Piezo2* in the distal colon. Arrows point to the triple-positive cells. Images are representative from three mice. **(b)** smRNA-FISH of *Piezo2/Piezo1/Tph1* in the indicated regions of SI. White arrowheads point to the absence of *Piezo2* in *Tph1+* cells in the SI. Yellow arrowheads point to sparse staining of *Piezo2* in the cells within the lamina propria. Images are representative from four mice. **(c,d)** qPCR validation of regional enriched genes in EC cells. GFP^−^ and GFP^+^ cells were sorted from the duodenum, jejunum, proximal and distal colon in the *Tph1-bacTRAP* mice. Relative gene expression was computed relative to the values in the GFP^−^ cells isolated from the duodenum (indicated as 1), after normalization by the aggregates of three house-keeping genes (*B2m, Gapdh, Rpl13a*). (c) *Tph1* and *Chga* were highly enriched in the GFP*+* cells from all regions. (d) *Trpa1*,*Trpm2,Cartpt and Ucn3* were highly enriched in small intestine. *** in (c) p<0.001; unpaired two-tailed Student’s *t*-test (between the GFP- and GFP+ cells). *** in (d) p<0.0001; two-way ANOVA test for both variables of GFP positivity and regions of gut, as well as the interaction of the two variables. Data are representative from three independent experiments. **(e,f)** smRNA-FISH assay for Htr4 and Tph1 in the distal colon. (e) Luminal side of the crypts is at the top of the image. An arrow points to the basal extension of a representative *Tph1^+^* cell projecting toward *Htr4+* epithelial cells (f). Image are representative from three wild-type mice. Scale bars in panels a,b,f: 10 µm; in panel e: 50 µm.

**Supplementary Figure 7.**
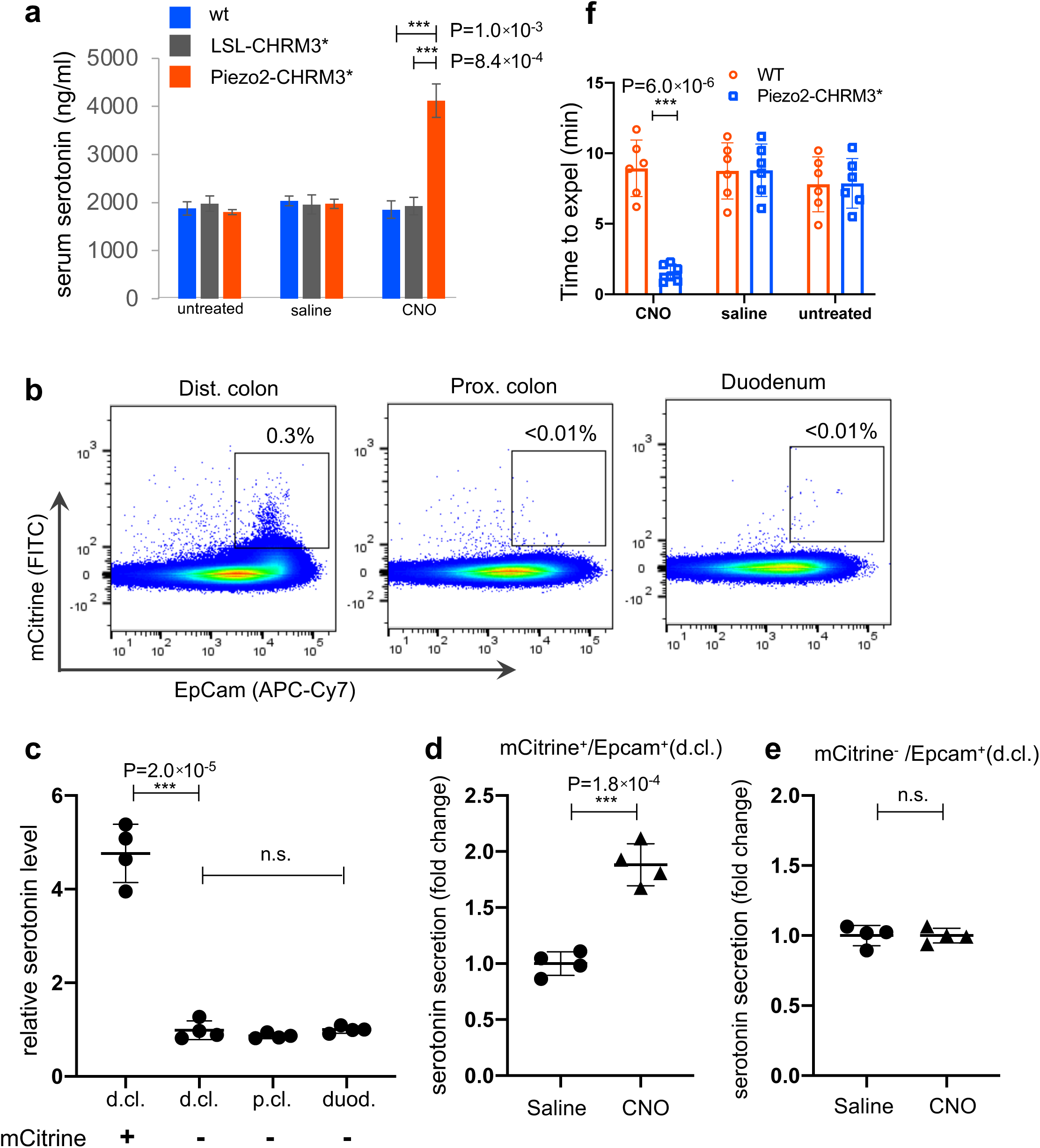
Chemogenetic activation of Piezo2 cells leads to serotonin release and accelerated colon motility. **(a)** Serum serotonin level examined 15 minutes after CNO administration (60 ng/kg) by ELISA. n=3 mice per group. **(b)** Representative flow cytometry results of dissociated epithelial cells from the indicated regions of the gut in the *Piezo2-CHRM3*/mCitrine* line. The Rosa26-LSL-CHRM3*/mCitrine locus is activated in a Piezo2-cre-specific manner. Percentages within indicated gates are shown. EpCam antibody was used to identify epithelial cells. Note that mCitrine^+^/EpCam^+^ cells were only observed in the distal colon of the *Piezo2-CHRM3*/mCitrine* mice. **(c)** Total serotonin levels shown for the sorted mCitrine^+^/EpCam^+^ cells from the distal colon (d.cl.), or mCitrine^−^/EpCam^+^ cells from the distal colon, proximal colon (p.cl.) and duodenum (duod.). Total serotonin was calculated as the sum of secreted serotonin and lysate serotonin, which was then normalized to the sum observed in the duodenal mCitrine^−^/EpCam^+^ cells. **(d,e)** Elevated serotonin secretion as observed in response to CNO treatment (10 ng/µl) in the mCitrine^+^/EpCam^+^ cells (d) but not the mCitrine^−^/EpCam^+^ cells isolated from the distal colon (e). Percentage of secreted serotonin/total serotonin was normalized to that of saline-treated cells. Serotonin level was assessed by ELISA. Data in (b-e) are based on four mice from two independent experiments. **(f)** BEA performed 20 minutes after CNO administration. n=6 mice per group.

**Supplementary Figure 8.**
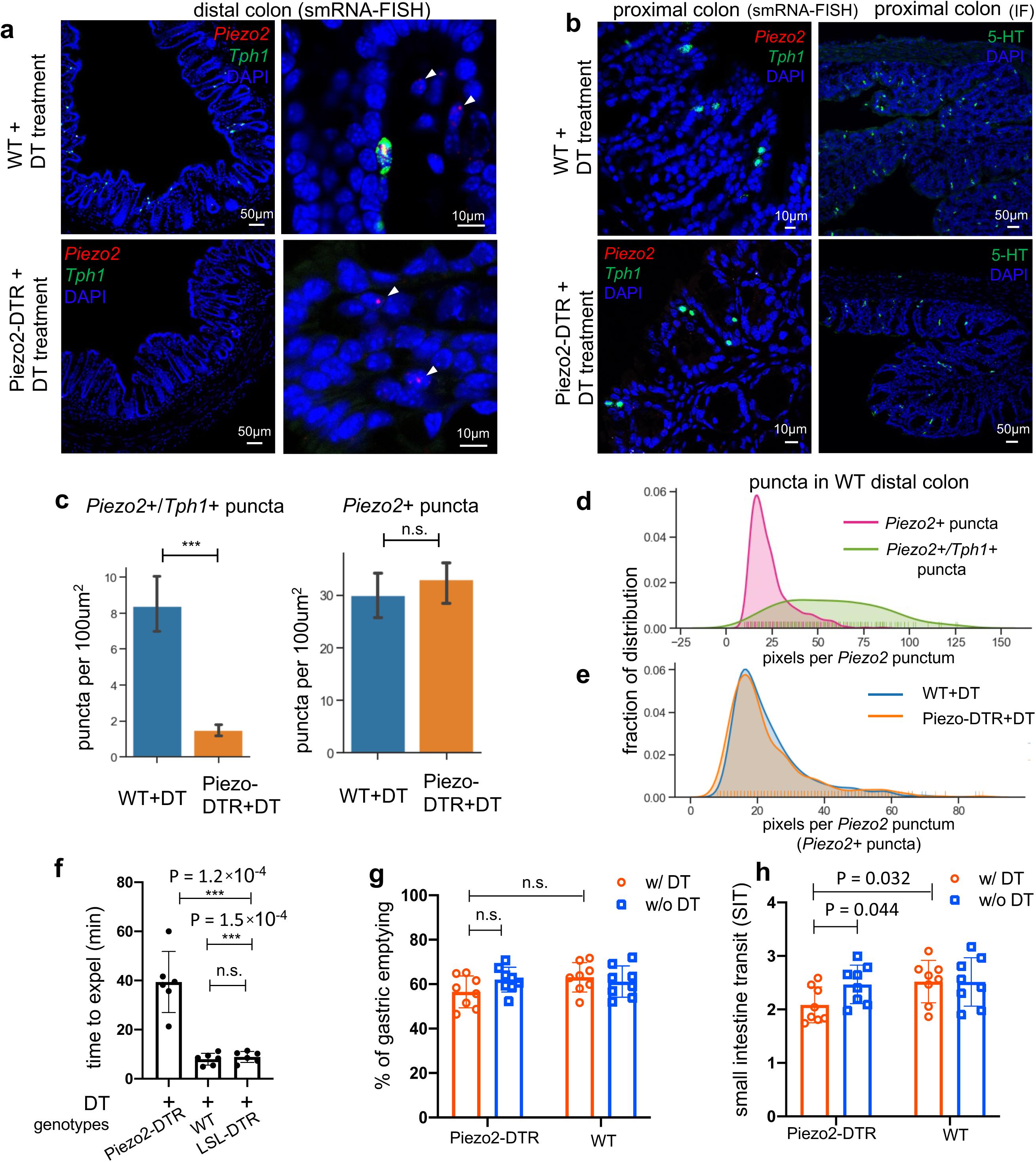
*Piezo2^+^/Ascl1^+^/Tph1^+^* cells required for normal colon motility. **(a)** *Piezo2* and *Tph1* transcripts determined by smRNA-FISH in the distal colon of wild type (WT, upper) or *Piezo2-DTR* (lower) mice after DT administration. Arrowheads point to the remaining submucosal staining of *Piezo2* in both WT and *Piezo2-DTR* mice. Images are representative from five animals per group. Scale bars: 10 µm or 50 µm, as indicated. **(b)** smRNA-FISH of Piezo2/Tph1 (left) and IF staining for 5-HT (right) in the proximal colon of the WT (upper) or *Piezo2-DTR* (lower) mice after DT administration. Data are representative from five animals per group. Scale bars: 10 µm or 50 µm, as indicated. **(c)** Signals of *Piezo2* single positivity (mostly in lamina propria) or *Tph1/Piezo2* double positivity (in the epithelial layer) were identified by smRNA-FISH in the distal colon of WT (blue) or *Piezo2-DTR* (orange) mice after DT administration and quantitated using a custom pipeline in Cellprofiler. The numbers of *Tph1/Piezo2* double-positive puncta (right) and *Piezo2* single-positive puncta (left) were based on eight mice from three independent experiments (4 of Piezo2-DTR+DT and 4 of WT+DT). n.s.: not significant, *** p<0.001; unpaired two-tailed Student’s *t*-test. **(d)** Distribution of pixels per punctum in the *Piezo2* channel quantitated in *Piezo2* single-positive puncta (pink) versus *Piezo2/Tph1* double-positive puncta (green). Pixels per punctum indicates expression levels of examined genes. **(e)** Distribution of pixels per punctum quantitated in *Piezo2* single-positive puncta identified in WT+DT (blue) versus Piezo2-DTR+DT (orange). **(f)** BEA performed after five days of consecutive treatment of DT in indicated groups. n=6 in each group. WT mice and mice with only LSL-DTR transgene were included as controls. All animals received the same DT treatment. **(g,h)** Gastric emptying and (g) small intestine transit (SIT) time (h). Animals were orally gavaged with methylcellulose supplemented with rhodamine B dextran (10 mg/ml). 15 minutes after gavage, the remaining rhodamine B dextran was determined in stomach and segments of intestine to assess upper GI motility. SIT was estimated by the position of the geometric center of the rhodamine B dextran in the small bowel. The geometric center values are distributed between 1 (minimal motility) and 10 (maximal motility). n=8 in each group. Scale bars: 10 µm or as indicated otherwise. Error bars denote standard deviation of the mean. * p<0.05, ** p<0.01, *** p<0.001; unpaired two-tailed Student’s *t*-test.

**Supplementary Figure 9.**
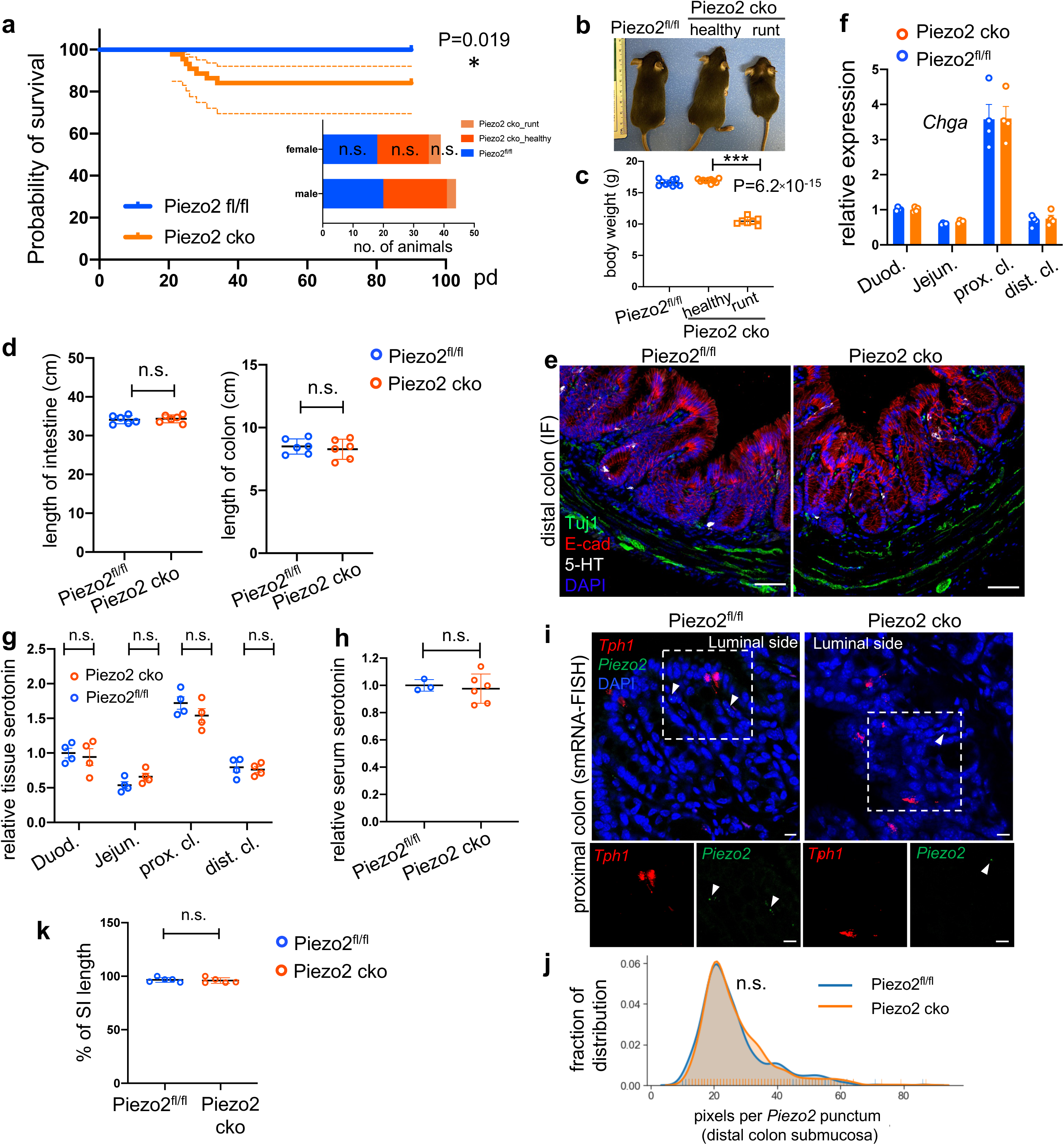
Epithelium *Piezo2* is required for efficient colon motility. **(a)** Kaplan-Meier survival curve of *Vilin-cre;Piezo2^fl/fl^* (Piezo2 cko, orange, n=44) in comparison to littermate controls of *Piezo2^fl/fl^*(blue, n=36). P=0.019, Mantel-Cox test. Dashed lines represent 95% confidence interval (CI). Inset: sex breakdown of the early mortality phenotype. n.s., not significant; Chi-square test. **(b)** Representative image of littermates (all male) of indicated genotypes before humane euthanasia of Piezo2 cko-runt animals. **(c)** Summary of body weights before humane euthanasia of the Piezo2 cko-runt animals. Both sexes are included. Each circle represents a datapoint. **(d)** Length of intestine (*left*) and colon (*right*) of the indicated animals at 8-weeks of age. Both sexes are included. Each circle represents a datapoint. **(e)** IF staining of 5-HT, E-cadherin and Tuj1 in the distal colon of *Piezo2^fl/fl^* (*left*) and Piezo2 cko (*right*) mice. Images are representative of three animals per group. **(f)** qPCR analysis of *Chga* in Piezo2 cko and *Piezo2^fl/fl^* epithelium. Gene expression was computed relative to the values in the *Piezo2^fl/fl^* duodenum, after normalization by the aggregates of three house-keeping genes (*B2m, Gapdh, Rpl13a*). Each circle represents one animal. n=4 per group. **(g)** Tissue serotonin level determined by ELISA and normalized by the serotonin level detected in the duodenum of the *Piezo2^fl/fl^* mice. n=4 for both Piezo2 cko (Vilin-cre;Piezo2^fl/fl^) and *Piezo2^fl/fl^*mice. n=4 per group. **(h)** Serum serotonin level determined by ELISA and normalized to the levels observed in *Piezo2^fl/fl^* mice. n=6 for Piezo2 cko and 3 for Piezo2^fl/fl^ mice. **(i)** smRNA-FISH of *Piezo2* and *Tph1* in the proximal colon of Piezo2^fl/fl^ (left) or *Piezo2 cko* (right) mice. Data are representative of three animals per group. **(j)** Distribution of pixels per punctum quantitated in *Piezo2* single-positive puncta identified in *Piezo2^fl/fl^*(blue) or Piezo2 cko (orange) distal colon. Pixels per punctum indicates expression levels of examined genes. **(k)** Summary data of fluorescent dye transit at 70 min after gavage. % of CI length = dye travel distance in SI ÷ full length of SI ×100%. Each circle represents one animal. Representative data from two independent experiments. Error bars stand for standard deviation of the mean. *p<0.05, ** p<0.01, *** p<0.001; unpaired two-tailed Student’s *t*-test.

**Supplementary Figure 10.**
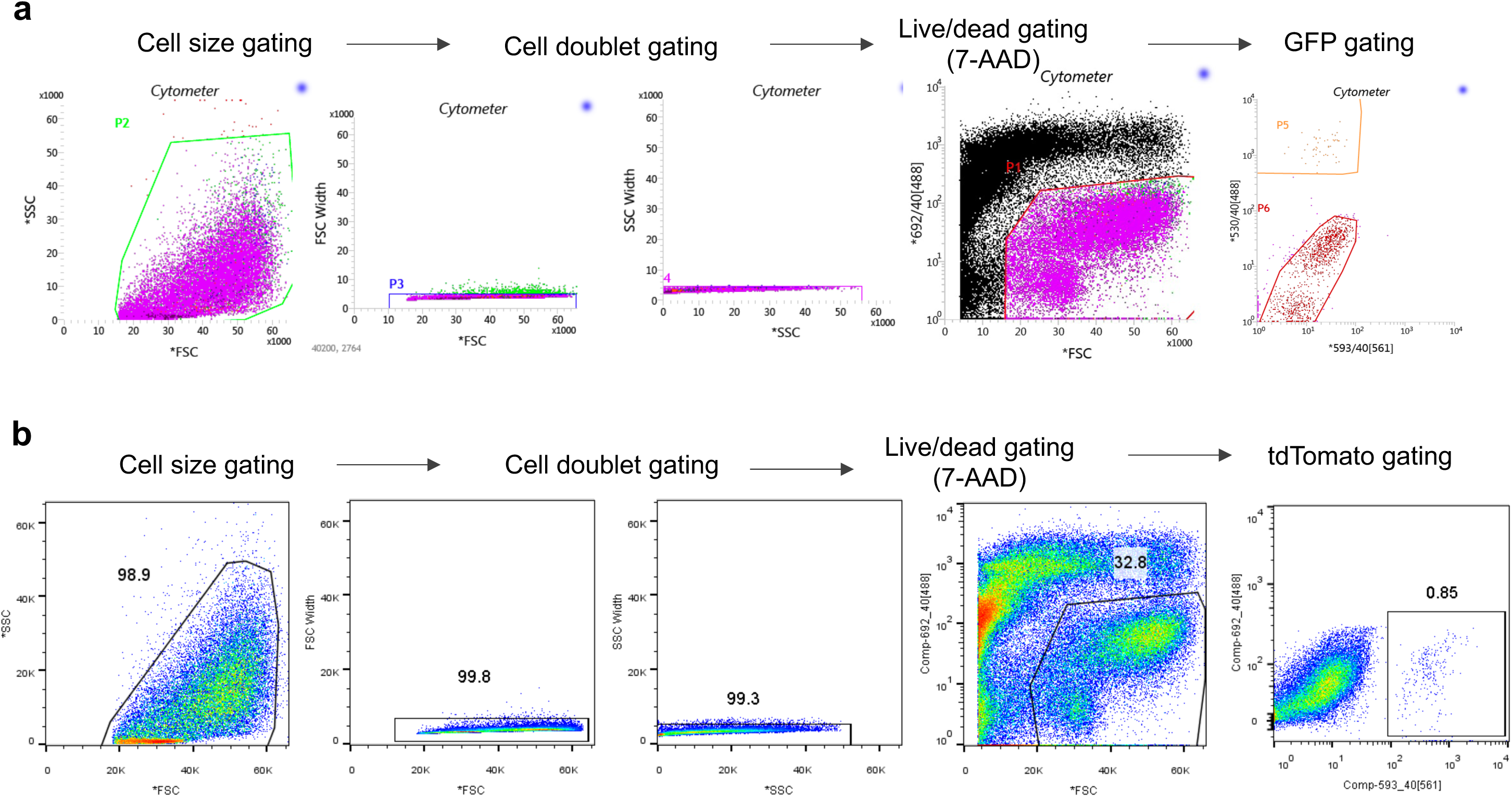
Examples of the FACS gating strategy. **(a)** FACS gating for GFP^+^ cells isolated from *Tph1-bacTRAP* mice. **(b)** FACS gating for tdTomato^+^ cells isolated from *Neurod1-tdTomato* mice.

## SUPPLEMENTARY TABLES

## Notes

### Competing Interest Statement

J.H. and J.W. are paid employees of Takeda Pharmaceuticals.

### Summary of Updates

1.Relevant to Fig. 1 and Suppl. Fig 1: Clarified that (i) some GFP is expressed in cells that have not yet fully committed to the EC lineage, or that there is some expression in cells outside this lineage, for example, in mast cells; (ii) it It is possible that the stem cell and transit amplifying cell clusters include cells that are in the process of differentiating into EC cells; and (iii) OSR2 and HOXB13 were preferentially enriched in the ileum and rectum, respectively, in the human samples. 2.Relevant to Fig. 4 and Suppl. Fig. 5: Clarified that our conclusion regarding the presence of subpopulations of EC cells in the proximal colon associated with different physiological roles is based on smFISH data alone. 3.Relevant to Fig. 6: Changed (quote) systematic administration (unquote) to (quote) systemic administration (unquote), (ii) discussed the reason for delayed small intestinal transit in the DTR experiments; (iii) added a comment speculating on why we did not see similar slowing of small intestinal transit in the Villlin-Cre Piezo2 KO; and (iv) added a comment on neural Piezo2 in the discussion, with relevant citations. 4.Throughout the manuscript, we rephrased (quote) nutrient sensing (unquote) to (quote) nutrient sensing and homeostasis (unquote) where appropriate.

