## Supplemental Table 1 for "Stratification of enterochromaffin cells by single-cell expression analysis"

**Supplementary Table 1: Summary of EC clusters and their potential physiological roles.**

| Cluster Number | Topological location | Molecular Identifiers | Distinguishing Genes | Possible Functions |
|---|---|---|---|---|
| 1, 7 | Duodenum(1) Jejunum (7) | *Tac1/Tph1* | *Neurog3, Neurod2, Tac1* | Precursors to other SI EC clusters |
| 2 | Duodenum villus | *Sct/Asic5/Tph1* | *Sct* (high), *Asic5 (low),Foxq1* | A major function may be to release secretin and 5-HT (both protecting against duodenal acidification) |
| 3, 10 | Duodenum villus (3) Jejunum villus (10) | *Trpm2/Cartpt/Tph1* | *Trpm2, Cartpt, Serpina1e, Tgfb1* | unknown |
| 4, 8 | Duodenum crypt (4) Jejunum crypt (8) | *Reg4/Tph1* | *Reg4, Ucn3,* | unknown |
| 5, 9 | Duodenum crypt (5) Jejunum crypt (9) | *Trpa1/Ucn3/Tph1* | *Trpa1,Ucn3, Reg4, Gstk1, Ces3a, Alb* | Hormone release in response to nutrients and phytochemicals, probably causing digestive enzyme release |
| 6 | Duodenum villus | *Cck/Oc3/Tph1* | *Cck, Ghrl, Oc3, Pzp, Cyp2j5, Habp2, Mc4r, Casr,*<br>*Crp, Tril, Tlr2, Tlr5, Lyzl4 (Bcam is high in cluster 6, but also detected in other EC cells)* | Possibly releases 5-HT and other hormones in reaction to luminal pathogens and tissue challenge. 5-HT released from these cells may initiate nausea. May also response to nutrients. |
| 11 | Proximal colon | *Iapp/Cpb2/Tph1* | *Iapp, Cpb2, Serpine1, Npy1r, (Pikb, Pde10a, Plet1)* | Coagulation and fibrinolysis<br>Possibly other roles |

| | | | | |
|---|---|---|---|---|
| 12 | Proximal colon | *Olfr558/Olfr78/Il12a/Tph1* | *Il12a, Olfr558, Olfr78, Reg4 (low), Igfbp7* | Microbial metabolite-sensing |
| 13 | Distal colon | *Piezo2/Olfr78/Foxj1/Tph1* | *Piezo2, Olfr78, Foxj1, Ascl1, Hoxb13* | Mechanosensitive EC that are important in motility control. They have basal long processes and primary cilia. May also respond to SCFA. |
| 14 | Distal colon | *Piezo2/Ascl1/Tph1* | *Piezo2, Ascl1, Hoxb13, Gper1, Vipr2* | Mechanosensitive EC that are important in motility control. They have basal long processes. |
