## Supplemental Table 4 for "Stratification of enterochromaffin cells by single-cell expression analysis"

**Supplementary Table 4. qPCR primers.**

| | |
|---|---|
| *Foxj1* | 5’ AGCCCAGAAGACTGGGAACT 3’ |
| | 5’ AATCCTTGGGCTTGAGGGAAC 3’ |
| *Ascl1* | 5’ GAATGGACTTTGGAAGCAGGATG 3’ |
| | 5’ TGCCCCTGTAGGTTGGCTG 3’ |
| *Piezo2-a* | 5’ GCACTCTACCTCAGGAAGACTG 3’ |
| | 5’ CAAAGCTGTGCCACCAGGTTCT 3’ |
| *Piezo2-b* | 5’ TCAAACACGCCAGTGACAAT 3’ |
| | 5’ TGTCTCTGAACAAAATGATGGTGA 3’ |
| *Trpa1* | 5’ GAGGATTGCTATGCAGGTGGA 3’ |
| | 5’ CGTGCCTGGGTCTATTTGGA 3’ |
| *Chga* | 5’ CCAAGGTGATGAAGTGCGTC 3’ |
| | 5’ GGTGTCGCAGGATAGAGAGGA 3’ |
| *Tph1* | 5’ TGTTGACTGCGACATCAGCCGA 3’ |
| | 5’ GGAAACCAAGGGACAGTCTCCA 3’ |
| *Trpm2* | 5’ AAGGATGTGGCTCTCACAGAC 3’ |
| | 5’ CGGGAACCCATACTCGACC 3’ |
| *B2m* | 5’ CACTGAATTCACCCCCACTGA 3’ |
| | 5’ TGTCTCGATCCCAGTAGACGG 3’ |
| *Rpl13a* | 5’ AGCAGATCTTGAGGTTACGGA 3’ |
| | 5’ GGAGTCCGTTGGTCTTGAGG 3’ |
| *Gapdh* | 5’ CTGGAGAAACCTGCCAAGTATG 3’ |
| | 5’ AGAGTGGGAGTTGCTGTTGAAG 3’ |
| *Cartpt* | 5’ AAGAAGTACGGCCAAGTCCC 3’ |
| | 5’ CAGTCACACAGCTTCCCGAT 3’ |
| *Ucn3* | 5’ AAGGCCAAGAATTTGCGAGC 3’ |
| | 5’ TGTCTTGATGTGCCACCCTC 3’ |
| *Il12a* | 5’ CCACTGGAACTACACAAGAACG 3’ |
| | 5’ ATGCTACCAAGGCACAGGGT 3’ |
