## Supplemental Table 2 for "Stratification of enterochromaffin cells by single-cell expression analysis"

**Supplementary Table 2: Primary and secondary antibodies used in this study.**

| Target | Name of Antibody | Source and reference to characterisation | Species raised in; clonality | Dilution used |
|---|---|---|---|---|
| 5-HT | #20079 | Incstar, (Hudson, WI, USA) (Cho et al. 2014) | Goat; polyclonal | 1:10000 |
| 5-HT | #20080 | ImmunoStar, (Hudson, WI, USA) | Rabbit; polyclonal | 1:1000 |
| Secretin | #H-067-04 | Phoenix Pharmaceuticals, (Mannheim, B-W, Germany) (Roy et al. 2012) | Rabbit; polyclonal | 1:2000 |
| CCK/gastrin | 28.2 | Gift from JH Walsh CURE/UCLA Antibody Core (Kovacs et al. 1989) | Mouse; monoclonal | 1:2000 |
| CCK/gastrin | 8007/5 | Rehfeld | Rabbit; | 1:8000 |
| GIP | Y-20 (SC-23554) | Santa Cruz Biotechnology, (Dallas, TX, USA) | Goat; polyclonal | 1:500 |
| Ghrelin | MF1601 | Shin YK et al, 2010 | Rabbit; polyclonal | 1:10000 |
| GFP | AB13970 | Abcam, (Melbourne, Australia) | Chicken; polyclonal | 1:2000 |
| Oxyntomodulin | AB-323-AO010 | Anshlabs (Webster, Texas, USA) | Mouse; monoclonal | 1:2000 |
| Substance P | SK1 | Eskay RL et al, 1980 | Rabbit; polyclonal | 1:1600 |
| Neurokinin A | 93K-6 | Too HP et al, 1990 | Rabbit; polyclonal | 1:400 |
| Neurotensin | SC-7592 | Santa Cruz Biotechnology, (Dallas, TX, USA) | Goat; polyclonal | 1:50 |

| | | | | |
|---|---|---|---|---|
| IFT88 | 13967-1-AP | Proteintech | Rb | 1:500 |
| Tuj1 | Ab18207 | Abcam | Rb | 1:1000 |
| E-cadherin | Ab11512 | Abcam | Rat | 1:1000 |
| Dclk1 | Ab31704 | Abcam | Rb | 1:1000 |
| Mouse IgG | 21202, Alexa Fluor 488 | Molecular Probes (Mulgrave, VIC, Australia) | Donkey; polyclonal | 1:500 |
| Chicken IgG | 103-545-155, Alexa Fluor 488 | Jackson ImmunoResearch (Waterford, QLD, Australia) | Goat; polyclonal | 1:500 |
| Rabbit IgG | A10042, Alexa Fluor 568 | Molecular Probes | Donkey; polyclonal | 1:800 |
| Sheep IgG | A-11015, Alexa Fluor 488 | Molecular Probes | Donkey; polyclonal | 1:500 |
| Sheep IgG | A21448, Alexa Fluor 647 | Molecular Probes | Donkey; polyclonal | 1:500 |
| Goat IgG | A21432, Alexa Fluor 555 | Molecular Probes | Donkey; polyclonal | 1:800 |
