## Supplemental Table 3 for "Stratification of enterochromaffin cells by single-cell expression analysis"

**Supplementary Table 3. Probes used for smRNA-FISH.**

| Target gene | Source | Cat# |
|---|---|---|
| Mm-*Tph1* | ACDBio/Bio-Techne | 318701-C2 |
| Mm-*Piezo2* | ACDBio/Bio-Techne | 400191-C3 |
| Mm-*Gpx3* | ACDBio/Bio-Techne | 400191-C3 |
| Mm-*Cartpt* | ACDBio/Bio-Techne | 432001 |
| Mm-*Stra6* | ACDBio/Bio-Techne | 450321 |
| Mm-*Cck* | ACDBio/Bio-Techne | 402271-C3 |
| Mm-*Foxj1* | ACDBio/Bio-Techne | 317091 |
| Mm-*Dnah9* | ACDBio/Bio-Techne | 556771-C3 |
| Mm-*Mc4r* | ACDBio/Bio-Techne | 402741 |
| Mm-*Casr* | ACDBio/Bio-Techne | 423451 |
| Mm-*Asic5* | ACDBio/Bio-Techne | 588601-C3 |
| Mm-*Pyy* | ACDBio/Bio-Techne | 420681-C3 |
| Mm-*Gpbar1* | ACDBio/Bio-Techne | 318451 |
| Mm-*Trpm2* | ACDBio/Bio-Techne | 313291 |
| Mm-*Ascl1* | ACDBio/Bio-Techne | 313291 |
| Mm-*Olfr558* | ACDBio/Bio-Techne | 316131-C2 |
| Mm-*Htr4* | ACDBio/Bio-Techne | 408241 |
| Mm-*Piezo1* | ACDBio/Bio-Techne | 500511 |
| Mm-*Onecut3* | ACDBio/Bio-Techne | 583241 |
| Mm-*Crp* | ACDBio/Bio-Techne | 583251 |
| Mm-*Cpb2* | ACDBio/Bio-Techne | 583261 |
| Mm-*Tlr2* | ACDBio/Bio-Techne | 317521 |
| Mm-*Tlr5* | ACDBio/Bio-Techne | 451601 |
| Mm-*F5* | ACDBio/Bio-Techne | 502411 |
| Mm-*Gip* | ACDBio/Bio-Techne | 451601 |
| Mm-*Cnr1* | ACDBio/Bio-Techne | 420721-C2 |
| Mm-*Ucn3* | ACDBio/Bio-Techne | 464861 |
| Mm-*Il12a* | ACDBio/Bio-Techne | 414881 |
| Mm-*Iapp* | ACDBio/Bio-Techne | 512571-C2 |
| Mm-*Olfr78* | ACDBio/Bio-Techne | 436601 |
